# A near-term iterative forecasting system successfully predicts reservoir hydrodynamics and partitions uncertainty in real time

**DOI:** 10.1101/2020.01.22.915538

**Authors:** R. Quinn Thomas, Renato J. Figueiredo, Vahid Daneshmand, Bethany J. Bookout, Laura K. Puckett, Cayelan C. Carey

**Affiliations:** Department of Forest Resources and Environmental Conservation, Virginia Tech, Blacksburg, VA, 24061, USA; Department of Electrical and Computer Engineering, University of Florida, Gainesville, FL, 32611, USA; Department of Biological Sciences, Virginia Tech, Blacksburg, VA, 24061, USA

**Keywords:** Data assimilation, Ecological forecasting, Ensemble Kalman filter, FLARE, General Lake Model, Water temperature

## Abstract

Freshwater ecosystems are experiencing greater variability due to human activities, necessitating new tools to anticipate future water quality. In response, we developed and deployed a real-time iterative water temperature forecasting system (FLARE – Forecasting Lake And Reservoir Ecosystems). FLARE is composed of: water quality and meteorology sensors that wirelessly stream data, a data assimilation algorithm that uses sensor observations to update predictions from a hydrodynamic model and calibrate model parameters, and an ensemble-based forecasting algorithm to generate forecasts that include uncertainty. Importantly, FLARE quantifies the contribution of different sources of uncertainty (driver data, initial conditions, model process, and parameters) to each daily forecast of water temperature at multiple depths. We applied FLARE to Falling Creek Reservoir (Vinton, Virginia, USA), a drinking water supply, during a 475-day period encompassing stratified and mixed thermal conditions. Aggregated across this period, root mean squared error (RMSE) of daily forecasted water temperatures was 1.13°C at the reservoir’s near-surface (1.0 m) for 7-day ahead forecasts and 1.62°C for 16-day ahead forecasts. The RMSE of forecasted water temperatures at the near-sediments (8.0 m) was 0.87°C for 7-day forecasts and 1.20°C for 16-day forecasts. FLARE successfully predicted the onset of fall turnover 4-14 days in advance in two sequential years. Uncertainty partitioning identified meteorology driver data as the dominant source of uncertainty in forecasts for most depths and thermal conditions, except for the near-sediments in summer, when model process uncertainty dominated. Overall, FLARE provides an open-source system for lake and reservoir water quality forecasting to improve real-time management.

**Key Points:**

- We created a real-time iterative lake water temperature forecasting system that uses sensors, data assimilation, and hydrodynamic modeling
- Our water quality forecasting system quantifies uncertainty in each daily forecast and is open-source
- 16-day future forecasted temperatures were within 1.4°C of observations over 16 months in a reservoir case study

## 1 Introduction

As a result of human activities, ecosystems around the globe are increasingly changing [*Stocker et al.*, 2013; *Ummenhofer and Meehl,* 2017], making it challenging for resource managers to consistently provide vital ecosystem services [*West et al.*, 2009]. In particular, managers of freshwater ecosystems, which have been more degraded than any other ecosystem on the planet [*Millennium Ecosystem Assessment,* 2005], are seeking new tools to anticipate future change and ensure clean water for drinking, fisheries, irrigation, industry, and recreation [*Brookes et al.*, 2014].

In response to this need, near-term, real-time iterative ecological forecasting has emerged as a solution to provide stakeholders, managers, and policy-makers crucial information about future ecosystem conditions [*Clark et al.*, 2001; *Dietze et al.*, 2018; *Luo et al.*, 2011]. Here, we define a real-time iterative forecast as a prediction of future ecosystem states with quantified uncertainty, generated from models that can be constantly updated with new data as they become available [adapted from *Clark et al.*, 2001; *Dietze,* 2017a; *Dietze et al.*, 2018; *Luo et al.*, 2011]. Multiple statistical approaches, such as Bayesian state-space modeling, particle filters, and ensemble filters, are used to estimate and propagate the different sources of uncertainty that contribute to the total uncertainty in a forecast (e.g., driver data, initial conditions, model process, and parameters) [e.g., *Clark et al.*, 2008; *Dietze,* 2017a; *Ouellet-Proulx et al.*, 2017a]. Fully specifying all of these uncertainty sources provides both an assessment of confidence in a forecast for managers as they interpret the forecasts for decision-making, and valuable information for researchers about how to improve forecasts [*Berthet et al.*, 2016; *Dietze,* 2017a; *Morss et al.*, 2008].

Real-time forecasts of water temperature with fully-specified uncertainties are particularly valuable for managers that oversee drinking water supply lakes and reservoirs, as waterbody temperatures can be very dynamic due to meteorological forcing, management, and seasonality [e.g., *Klug et al.*, 2012; *Mi et al.*, 2019; *Schmidt et al.*, 2018; *Sharma et al.*, 2015]. Water temperature is closely related to many water quality metrics and defines the physical conditions for multiple ecological processes, including microbial and algal growth, dissolved oxygen saturation, the release of chemical constituents from sediments into the water column, and habitat suitability for organisms, such as fish [*Butcher et al.*, 2015; *Carey et al.*, 2012; *Delpla et al.*, 2009; *Jöhnk et al.,* 2008]. As a result, depth profiles of temperature data are used to determine withdrawal depths for water treatment, dam release schedules for hydropower generation and downstream flows, and lake and reservoir water quality management [*Çalişkan and Elçi,* 2008; *Casamitjana et al.*, 2003; *Hague and Patterson,* 2014; *Huang et al.*, 2011; *Pike et al.*, 2013; *Weber et al.*, 2017]. Water temperature depth profiles also determine the strength of thermal stratification, i.e., if there are discrete epilimnetic (surface) and hypolimnetic (bottom) layers or isothermal (fully-mixed) conditions [*Read et al.*, 2011]. When waterbodies transition from stratified to mixed conditions during the onset of fall turnover, nutrients and metals that have accumulated in the hypolimnion during the summer are mixed throughout the water column, decreasing water quality [*Cooke et al.*, 2005; *Effler and Matthews,* 2008]. Consequently, real-time iterative forecasts of water temperature profiles would allow managers to preemptively respond to impending poor water quality during fall turnover and other episodic events (e.g., storms) that alter water temperature and thermal stratification.

Here, we introduce a forecasting system (FLARE, Forecasting Lake And Reservoir Ecosystems) that generates automated 16-day water temperature forecasts in real time (Figure 1). FLARE is composed of: 1) water quality and meteorology sensors deployed in a lake or reservoir that wirelessly stream data, 2) a data assimilation algorithm that uses sensor observations to update water temperature predictions from a hydrodynamic model and to calibrate model parameters, and 3) an ensemble-based forecasting algorithm to generate forecasts that quantify the sources of forecast uncertainty. FLARE quantifies uncertainty from driver data (i.e., the uncertainty in future weather or inflow forecasts that are needed to run the hydrodynamic model), initial conditions (i.e., the uncertainty observed in water temperatures on the first day of the forecast), model process (i.e., the capacity of a calibrated model to reproduce observations), and model parameters [following *Dietze,* 2017a]. The forecasting system samples from these sources of uncertainty to generate probability distributions for water temperature at multiple depths and can generate probability distributions of hydrodynamic events such as the occurrence of fall turnover.

**Figure 1.**
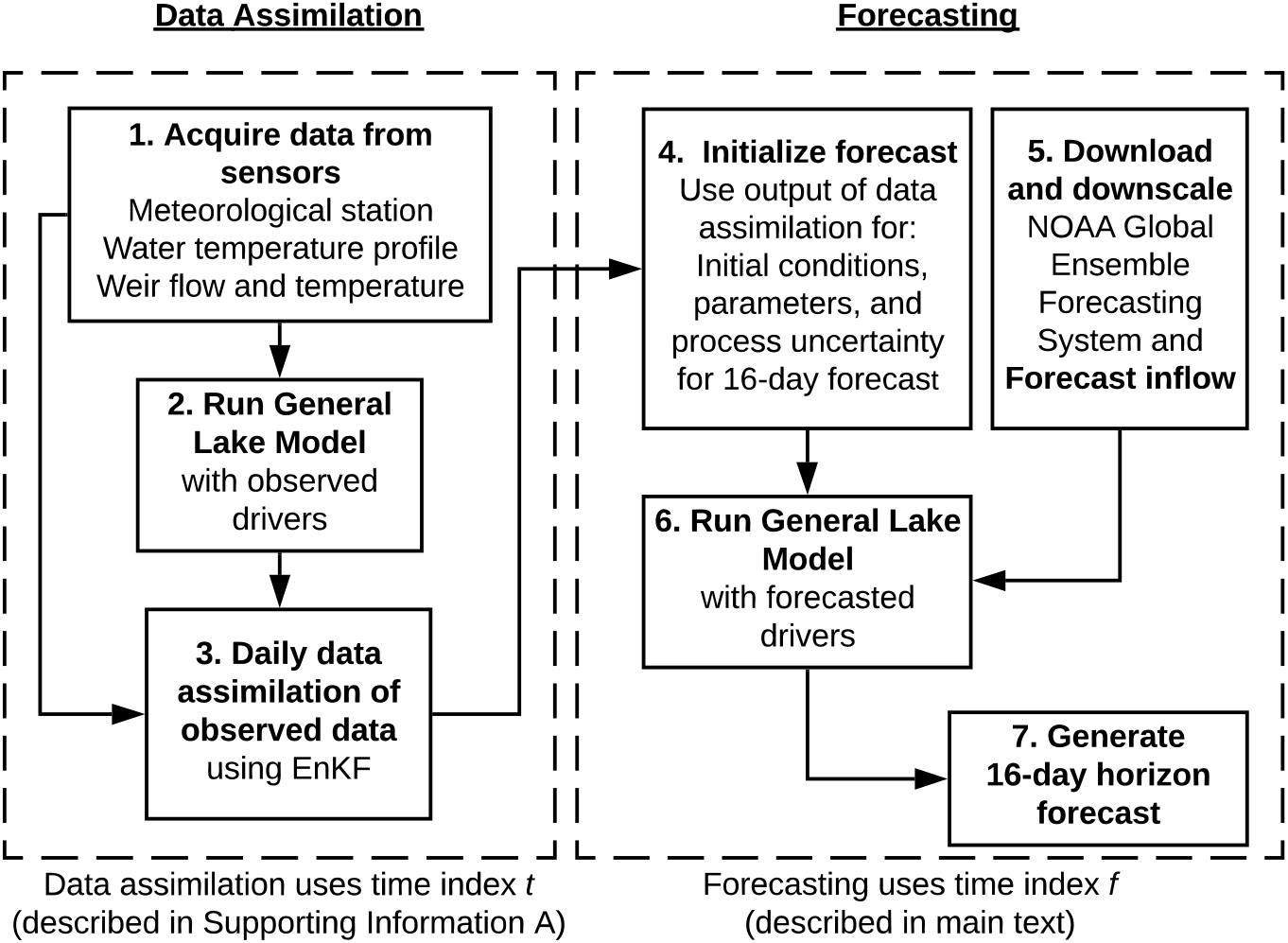
A simplified conceptual diagram describing the major data assimilation and forecasting steps in the daily iterative forecasting cycle for the FLARE (Forecasting Lake And Reservoir Ecosystems) forecasting system. The forecasting cycle has two key stages in which data are assimilated each day (left box) to produce the initial conditions for a 16-day forecast (right box). EnKF = Ensemble Kalman Filter; NOAA = National Oceanic and Atmospheric Administration.

FLARE builds off of previous studies that have developed forecasting systems with data assimilation to predict water temperature [e.g., *Bal et al.*, 2014; *Baracchini et al.*, 2020b; *Caissie et al.*, 2016; *Hague and Patterson,* 2014; *Huang et al.*, 2011; *Mestekemper et al.*, 2010; *Ouellet-Proulx et al.*, 2017a; *Pike et al.*, 2013] in four key ways. First, FLARE was built using all opensource software and model components and is completely automated via its cyberinfrastructure: no “human-in-the-loop” is needed to retrieve real-time observations for data assimilation, run models, or trigger any steps to generate daily forecasts. Second, unlike many real-time water forecasting systems that depend on governmental agencies for the collection of data used for assimilation [e.g., *Ouellet-Proulx et al.*, 2017a; *Pike et al.*, 2013], FLARE forecasts can be run and updated with water temperature data commonly collected by local water managers. These features enable scalability to a range of waterbodies without existing infrastructure. Third, FLARE daily forecasts extend up to 16 days in the future, which is a longer forecast horizon than existing water temperature forecasting systems, which have forecast horizons ranging from 3 days [*Caissie et al.*, 2016] to 10 days [*Hague and Patterson,* 2014]. Fourth, FLARE propagates and partitions multiple sources of forecast uncertainty, which are updated over time, allowing for the evaluation of both forecast accuracy and the reliability of uncertainty estimation. Despite the importance of quantifying multiple uncertainty sources, few water resource forecasting studies quantify more than one or two sources of uncertainty and when they do, they typically only include initial conditions uncertainty (via data assimilation) and meteorological uncertainty (via ensemble weather forecasts) [e.g., *Baracchini et al.*, 2020b; *Komatsu et al.*, 2007; *Ouellet-Proulx et al.*, 2017a; *Ouellet-Proulx et al.*, 2017b; *Page et al.*, 2018]. Furthermore, they rarely partition the relative contributions of the individual sources of uncertainty to the total forecast uncertainty [but see *Ouellet-Proulx et al.*, 2017a].

We set up FLARE to generate automated, probabilistic water temperature forecasts for a drinking water reservoir over 16 months (475 days) to address the following questions: 1) How does forecasting performance, as evaluated against observations and a null persistence model, differ between thermally-stratified vs. mixed conditions, two key regimes of lake thermal dynamics?, 2) How well does the forecasting system predict the onset of fall turnover?, and 3) What are the contributions of different sources of uncertainty to the forecasts, and how do they vary between thermally-stratified vs. mixed conditions?

## 2 Methods

We developed a forecasting system (FLARE; Forecasting Lake And Reservoir Ecosystems) that predicts water temperature at any set of specified depths in a lake or reservoir for a 16-day time horizon using a physics-based hydrodynamic model (Figure 1). The system uses data assimilation of observed data to generate the initial conditions, parameters, and uncertainty estimates for a forecast that extends 16 days in the future. Each day, the data assimilation is advanced one daily time step before launching the forecast of future conditions. We describe the forecasting methods below and the data assimilation methods in Supporting Information A.

### 2.1 Hydrodynamic model

FLARE simulated reservoir hydrodynamics with the General Lake Model (GLM), a one-dimensional (1-D) vertical stratification model [*Hipsey et al.*, 2019]. We used GLM because: 1) the model has successfully reproduced observed water temperature profiles in lakes around the world with varying mixing regime, climate, and morphology [*Bruce et al.*, 2018]; 2) GLM is an open-source, community-developed model and thus scalable to other waterbodies; and 3) GLM has low computational needs, enabling many model ensemble members to be run quickly and efficiently, a requirement for real-time iterative forecasting. GLM runs at a daily time step with sub-daily (hourly) meteorological drivers and daily inflow drivers.

We set all but three highly-sensitive parameters equal to the values reported in the default GLM version 3.1.0a model [*Hipsey et al.*, 2019], and FLARE updated the highly-sensitive parameters at each model time step. The three parameters that had automated tuning using data assimilation were selected using a sensitivity analysis described in Supporting Information A and were: one parameter that scales the incoming shortwave radiation (sw_factor) and two parameters defining the sediment temperatures in the hypolimnion (zone1temp, 5 – 9.3 m) and epilimnion (zone2temp: 0 – 5 m). For driver data, GLM requires hourly meteorological data on downwelling shortwave radiation (Wm-^2^), downwelling longwave radiation (W m^-2^), air temperature (°C), wind speed (m s-1), relative humidity (%), and precipitation (m day^-1^) as well as daily rates of water inflow (m3 day^-1^) and inflow temperature *(°C)* and daily rates of outflow (m3 day^-1^). The full GLM configuration with model driver files is provided in Supporting Information B.

### 2.2 Ensemble forecasting approach

The FLARE system uses an ensemble approach to numerically simulate and propagate forecast uncertainty into the future. The ensemble forecasting is based on eqn. 1:

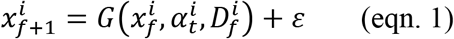

where G() is the GLM model that requires a vector of water temperatures at the modeled depths as initial conditions 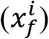, a vector of the three calibrated model parameters 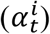, and a set of model drivers (*D*, which encompassed weather, inflows, and outflows). The index *i* in eqn. 1 represents the *i*th ensemble member and the subscript *f* is the day (0 to 16) in the 16-day forecast horizon. The index *t* is the number of days since the initiation of forecasting and data assimilation (i.e., 0 = the day the automated forecasting system is deployed). The index *t* is different from *f* because *t* increases each day throughout a multi-year forecasting period, while *f* is reset to zero each day before initiating a new 16-day forecast. *ε* represents the contribution of process uncertainty [i.e., errors due to an imperfect representation of physical processes in a model; *Dietze*, 2017a] to total forecast uncertainty and is randomly drawn from a multivariate normal distribution with a mean of zero and a covariance matrix of Σ_*t*_. The covariance matrix (Σ_*t*_) represents the uncertainty at each depth (the diagonals of Σ_*t*_) and the covariance of uncertainty between depths (the off-diagonals of *Σ_t_*). *Σ_t_* is calculated from the residuals between the ensemble mean and the observed temperatures before data are assimilated. The subscript *t* on Σ_*t*_ signifies that the covariance matrix is determined by data assimilation of historical observations and is constant over each day *f* in the 16-day forecast horizon. Eqn. 1 is applied at the daily time step and the process uncertainty is added each day of the forecast.

Each 16-day forecast requires initial conditions of the water temperature at each modeled depth (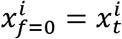, where *t* is the number of days since the automated forecasting system is deployed) and model parameters 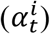 on day *f* = 0 of the forecast. The variance in *x_t_* and *α_t_* across ensemble members when the forecast is initialized represents the contribution of initial conditions and parameters, respectively, to the total forecast uncertainty. To generate these initial conditions while also calibrating the three focal parameters, FLARE assimilates temperature sensor observations from the previous day into the GLM using an Ensemble Kalman Filter [EnKF; *Evensen*, 2003; *Evensen*, 2009]. We used data assimilation rather than simply specifying the initial conditions from the observations because the EnKF: 1) enables the generation of initial conditions when sensor data were not available (e.g., during maintenance), 2) mechanistically interpolates water temperature between sensor depths, 3) enables the automated calibration of the three highly-sensitive model parameters, and 4) generates a historical data product of water temperature with spatial and temporal gap-filling. The EnKF method of data assimilation is well-suited for non-linear mechanistic models like the GLM and enables ensemble-based forecasts of future states [*Baracchini et al.*, 2020a; *Clark et al.*, 2008; *Dietze,* 2017a; *Page et al.*, 2018]. Our implementation of the EnKF with state augmentation to calibrate parameters (Supporting Information A) follows *Zhang et al.* [2017].

To quantify the contribution of future meteorological conditions to the total forecast uncertainty, each of the *i* water temperature ensemble members is assigned one of the ensemble meteorological forecast members from the National Oceanic and Atmospheric Administration Global Ensemble Forecasting System (NOAA GEFS) throughout the 16-day forecast horizon. NOAA GEFS provides ensemble forecasts with 21 members, with each member representing slightly modified versions of the model that result in different predictions of future weather conditions at a 16-day time horizon. Each day that a forecast is generated, the meteorological driver data required by the GLM from NOAA GEFS are downloaded for the 12:00:00 UTC forecast using the rNOMADS package (version 2.4.2) in R [*Bowman,* 2019].

Additionally, our forecasts include the uncertainty from statistically downscaling the gridded NOAA GEFS meteorological forecasts to local meteorological conditions. First, we downscaled the NOAA GEFS forecasts to a local site using a linear relationship between the NOAA GEFS forecast and observed meteorology. Next, we added random noise to each meteorological variable for each day of each NOAA GEFS ensemble member. This random noise was generated by a multivariate normal distribution that described the covariance in the relationship between the observed meteorology and NOAA GEFS forecasted meteorology for the meteorological variables required by GLM. By adding random error from this multivariate normal distribution to each member of the 21-member NOAA GEFS, we generated an ensemble of meteorological drivers that represented both NOAA GEFS forecast and downscaling uncertainty (see Supporting Information C for more detail about the downscaling methods).

In addition to meteorological driver uncertainty, we include driver uncertainty from future inflow discharge and temperature of the primary tributary to the lake or reservoir. For simplicity, we assume here that there is only one primary inflow, but additional inflows could be easily added in applications to other waterbodies. The inflow is forecasted using a simple first-order auto-regressive empirical model that predicts tomorrow’s inflow discharge 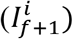 as function of today’s flow 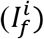, today’s daily precipitation 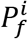, and random error *(ε)* that is normally distributed with a mean of 0 and a standard deviation of 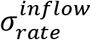:

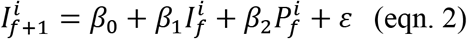

in which *β*_0_,*β*_1_, and *β*_2_ are regression coefficients. 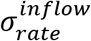 is estimated using the residuals from the regression fit. Eqn. 2 is iteratively applied through the 16-day forecast horizon for each downscaled meteorological driver ensemble member.

Similar to inflow, we forecast future inflow water temperature *(T)* as a function of today’s water temperature 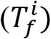, today’s air temperature 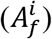, and random error *(ε)* that is normally distributed with a mean of 0 and a standard deviation of 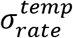

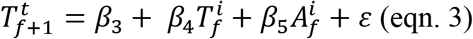

*β*_3_,*β*_4_, and *β*_5_ are regression coefficients. 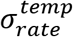 is estimated using the residuals from the regression fit. After inflow rate is calculated for each ensemble member, the outflow for that ensemble member is set to equal the inflow to maintain constant water levels.

### 2.3 Application of forecasting system

#### 2.3.1 Site description

We applied and evaluated FLARE at Falling Creek Reservoir (FCR), a dimictic, eutrophic reservoir located in Vinton, Virginia, USA (37.30°N,79.84°W). FCR is a shallow (maximum depth = 9.3 m, mean depth = 4 m), small (surface area = 0.119 km2) reservoir [*Gerling et al.*, 2016]. The lake generally exhibits summer thermal stratification from May to October and is ice-covered for short durations during December to March [*Carey,* 2019]. FCR is primarily fed by one upstream tributary and was maintained at full pond throughout this study by the Western Virginia Water Authority (WVWA), who own and manage the reservoir as a drinking water supply [*Gerling et al.*, 2014; *Gerling et al.*, 2016].

We used high-frequency sensors to monitor FCR water temperature [*Carey et al.*, 2020b] and meteorology [*Carey et al.*, 2020a], and measure the inflow discharge rate and water temperature of the primary tributary entering FCR through a weir [*Carey et al.*, 2020c]. Descriptions of the sensor array and methods for real-time wireless transfer of data to cloud storage are in Supporting Information D.

#### 2.3.2 Description of forecasting analysis

Our application of the forecasting system at FCR was from 11 July 2018 to 15 December 2019. There were no 16-day forecasts produced between 11 July 2018 and 27 August 2018 in order to use observed meteorology and data assimilation during this period to estimate the Σ_*t*_ process uncertainty covariance matrix and for the automated tuning of the three GLM parameters (see Supporting Information A). Starting on 28 August 2018, 16-day forecasts were produced each day until 15 December 2019. We forecasted water temperature on 0.33-m depth intervals starting at the surface through 9.0 m at the sediments. This resulted in 28 model states of water temperature depths in the *x* matrix (eqn. 1). We had sensor observations for 11 of the 28 modeled depths (0.1, 1.0, 1.6, 2.0, 3.0, 4.0, 5.0, 6.0, 7.0, 8.0, and 9.0 m; Supporting Information D).

In the data assimilation and forecasting steps of the daily workflow (Figure 1), we used N = 441 ensemble members. Even though fewer ensemble members have been shown to be effective when applying the EnKF in other studies [e.g., *Loos et al.*, 2020], our ensemble size was selected to adequately sample forecasted meteorology drivers to include uncertainty from both NOAA GEFS and its statistical downscaling. In the forecasting step for this application (Figure 1), we generated 21 members of our downscaling ensemble for each of the 21 NOAA GEF ensemble members, resulting in a total ensemble size of N = 441. The same ensemble size was applied to the data assimilation step to match the forecasting step.

We estimated the coefficients for the inflow discharge and temperature forecasting models in eqns. 2 and 3 using data from 1 January 2017 to 12 July 2018 (*β*_0_ = 0.001, *β*_1_= 0.948, *β*_2_ = 0.358, 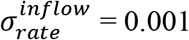, *β*_3_ = 0.203, *β*_4_= 0.942, *β*_5_ = 0.043, 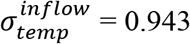). Both regressions had high R_2_ (eqn. 2: R_2_ = 0.98; eqn. 3: R_2_ = 0.93) and statistically significant parameters (all parameters p < 0.05). In the forecast, each of the 441 meteorological ensembles were used to separately calculate inflow discharge and temperature. Following eqns. 2 and 3, random noise was added to each ensemble member to represent uncertainty in forecasting inflow. We set the inflow discharge and temperature at the beginning of each 16-day forecast (*f*=0) directly from the sensors located on the primary tributary entering FCR.

Beginning 28 August 2018, 16-day horizon forecasts were produced each day until the end of the forecasting period on 15 December 2019. The 475 daily forecasts generated over this period included summer stratified, fall mixed, under-ice stratified, and spring mixed conditions in the reservoir [*Carey et al.*, 2020b]. Our iterative daily forecasting cycle (Figure 1) entailed: 1) retrieving the previous 24 hours of meteorological, inflow, and water temperature data from the reservoir sensor network, 2) advancing the model states (i.e., the water temperatures at the modeled depths) one day using the observed meteorology and inflows as drivers to the GLM model, 3) using the EnKF to assimilate observed water temperature and update states and parameters, and 4) initiating a 16-day forecast using updated states and parameters as initial conditions and parameters, which started at 12:00:00 UTC of the current day. As result of the daily data assimilation set in the forecasting cycle, parameter distributions and process uncertainty (Σ_*t*_) were updated through each iteration of the forecasting cycle. Once a 16-day forecast was launched from the initial conditions set by data assimilation, the parameters did not change over its 16-day horizon.

We also used a persistence null model to forecast water temperatures from 28 August 2018 to 15 December 2019 as a baseline for evaluating forecast skill. We used a persistence null model rather than a climatology null model because we lacked the multiple years of daily water temperature observations that would be necessary to generate historical averages. Our null model assumed water temperature did not change over the 16-day horizon based on eqn. 4.

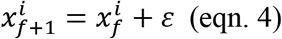

where *ε* is represents the contribution of process uncertainty to total forecast uncertainty and is randomly drawn from a multivariate normal distribution with a mean of zero and a covariance matrix of 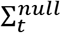. We applied the same data assimilation methods used to determine the initial conditions 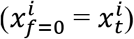 and process uncertainty covariance matrix 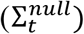 in the forecast (Supporting Information A) to the persistence null forecast.

#### 2.3.3 Evaluation of forecasts

We used multiple metrics to evaluate forecast performance. First, we calculated root mean squared error (RMSE) and the mean Continuous Ranked Probability Score (CRPS) for the forecasted water temperature (*F*) and the observed water temperature (*y*) at each day in the 16-day forecast horizon. CRPS is analogous to mean absolute error and is used with probability distributions produced by ensemble forecasts [*Gneiting et al.*, 2005]. CRPS is defined as:

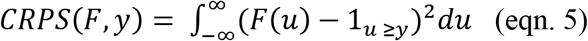

where 1_*u ≥y*_ is a step function that takes the value 1 if u ≥ y and the value 0 otherwise [*Gneiting et al.*, 2005]. We used the “verification” package in R [*Gilleland,* 2015] to calculate CRPS. The forecast skill was assessed by comparing the CRPS of the forecast (*CRPS_forecast_*) and the null model *(CRPS_null_*) using a skill score:

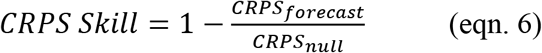

Thus, we report three CRPS values: *CRPS_forecast_*, *CRPS_null_*, and CRPS forecast skill, which was a normalized value in which zero indicated a forecast that performed the same as the null, one was a perfect forecast in that it predicted observations with no error, and values less than one indicated a forecast that performed worse than the null.

Second, bias was assessed using the absolute difference between the mean forecasted water temperature *(f_t_)* and the observed water temperature *(o_t_)* at each day in the 16-day forecast.

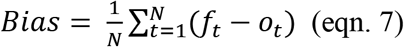

Third, the reliability of the confidence intervals was also assessed by calculating the total number of observations contained in a specific confidence interval [i.e., a reliability diagram; *Bröcker and Smith,* 2007]. We considered a forecast to be well-calibrated if its 90% confidence intervals contained 90% of the observations over the 100 days of forecasts. The confidence interval would be over-confident if fewer than 90% of observations were in this interval and would be under-confident if more than 90% observations were in this interval. We calculated the proportion of observations in the 10, 20, 30, 40, 50, 60, 70, 80, and 90th confidence intervals for 1-day, 7-day, and 16-day time horizons.

In our evaluation, we averaged RMSE, CRPS, bias, and reliability of confidence intervals for each day within the 16-day forecast horizon over the 475 daily forecasts to calculate aggregated forecast performance. In addition, we classified each of the 475 days as either thermally-stratified or mixed to compare relative forecast performance between the two thermal regimes. We applied the turnover criterion of *McClure et al.* [2018]: days with water temperatures at the near-surface (1.0 m) and near-sediments (8.0 m) at 12:00:00 UTC (the beginning of each forecast’s daily time step) that were <1°C different were categorized as mixed, and days with water temperatures at 1.0 m and 8.0 m that were ≥1°C different were categorized as thermally-stratified. Given that there was only intermittent ice cover on the reservoir for N = 11 days total during the forecasting period and stable inverse thermal stratification never occurred [*Carey,* 2019], days with ice were classified as mixed.

Finally, we evaluated the ability of the forecasting system to predict the day that fall turnover was first observed at the reservoir in both years. In the forecasts prior to turnover, we calculated the proportion of ensemble members that predicted the 1.0 m and 8.0 m water temperatures to be <1°C different on each day in the forecast. We expected this probability to increase for the day of turnover relative to other days as the onset of turnover approached.

#### 2.3.4 Partitioning of uncertainty

We isolated the contribution of each uncertainty source by removing the contributions of all other uncertainty sources in a set of scenarios run for each day over the forecasting period. Initial condition uncertainty was removed from the scenarios by initializing all ensemble members at *f* = 0 (eqn. 1) with the mean from all ensemble members. Parameter uncertainty was removed by assigning all 441 ensemble members to have the same ensemble means for each of the three automatically-calibrated parameters. Process uncertainty was removed by excluding the error term *(ε)* in eqn 1. NOAA GEFS meteorological driver data uncertainty was removed by using the ensemble mean from the 21-member NOAA GEFS ensemble instead of each individual NOAA GEFS ensemble member. Meteorology statistical downscaling uncertainty was removed by not sampling from the multivariate normal distribution describing the variance and covariance in downscaling error among meteorology variables. Inflow driver data uncertainty was removed by not sampling from the normally distributed uncertainty in eqns. 2 and 3. In our analysis, we compared the variance contributed by each isolated uncertainty source and summed the uncertainty sources to calculate the total forecast variance separately each day in the 16-day horizon. To examine how the uncertainty contributions varied over time, we partitioned the uncertainty of the forecasts generated on the first day of each month, resulting in 16 uncertainty-partitioned forecasts during 28 August 2018 to 15 December 2019. We separately averaged the partitioned uncertainty for the N = 9 forecasts generated during thermally-stratified conditions and N = 7 forecasts generated during mixed conditions.

## 3. Results

### 3.1 Observational water temperature data

Falling Creek Reservoir exhibited summer thermal stratification from the beginning of the monitoring period on 11 July 2018 until the onset of fall turnover occurred on 22 October 2018, and then remained mixed until thermal stratification began to set up on 12 March 2019 (Figure 2A). The reservoir remained thermally-stratified throughout the summer until the onset of the second fall turnover on 24 October 2019. We observed similar thermal dynamics between the two years: observed water temperatures at the reservoir’s near-surface (1.0 m) peaked on 17 July (28.1°C) in 2018 and on 21 July (29.3°C) in 2019. In both years, fall turnover was preceded by rapidly-cooling surface water temperatures, which decreased from 22.6 to 14.7°C at 1.0 m in the 14 days prior to 22 October 2018 and from 20.0 to 14.6°C in the 14 days prior to 24 October 2019. In the mixed periods of 2018 and 2019 (Figure 2B), the 1.0 and 8.0 m depth thermistors recorded water temperatures at 12:00:00 UTC that had a mean difference of 0.23°C (±0.55°C, 1 S.D).

**Figure 2.**
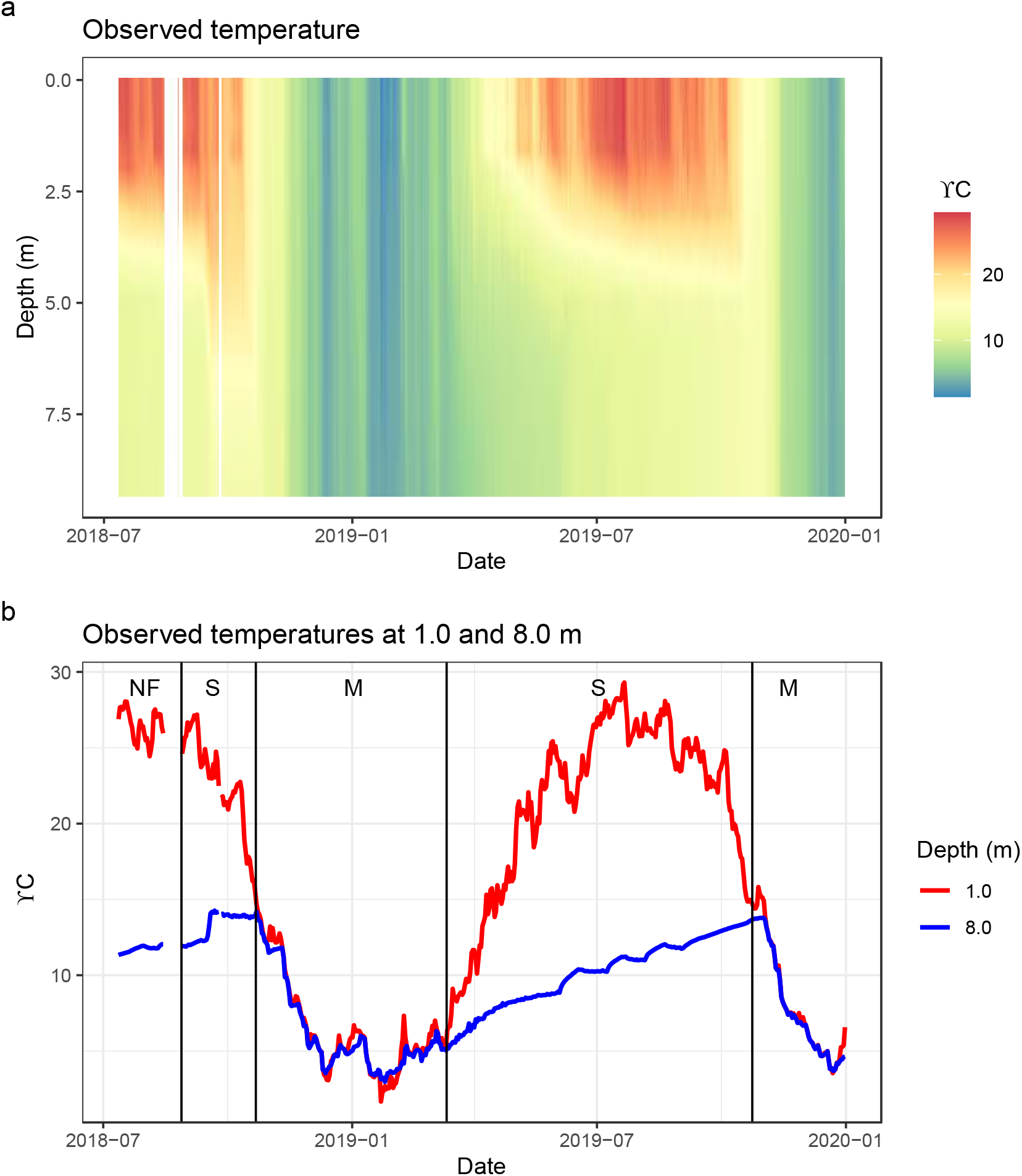
Observed water temperature depth profiles in Falling Creek Reservoir during the forecasting period (28 August 2018 to 15 December 2019). (a) A heat map showing the observed temperatures through the water column for the forecasting period based on water temperature sensor observations measured every 10 minutes at 0.1, 1.0, 1.6, 2.0, 3.0, 4.0. 5.0, 6.0, 7.0, 8.0, and 9.0 m depths. White values denote days with missing water temperature sensor data as a result of sensor maintenance. (b) A comparison of the observed water temperature for two depths (1.0 m: red, 8.0 m: blue). NF (Not Forecasted) denotes the period when data assimilation was performed but no forecasts were generated. The S and M labels refer to the Stratified and Mixed periods, respectively.

### 3.2 GLM model and ensemble Kalman filter (EnKF) performance

The GLM model was able to successfully reproduce observed patterns in reservoir water temperature following data assimilation. With daily assimilation of observed water temperatures using the EnKF and observed (not forecasted) meteorological and inflow drivers, the RMSE for the water temperature predicted by GLM at 1.0 m and 8.0 m depth was 0.06°C and 0.07°C, respectively, during the period when data assimilation occurred (11 July 2018 – 15 December 2019; Figure 3). Data assimilation substantially improved GLM predictions, as the RMSE for predicted water temperatures at 1.0 m and 8.0 m using observed drivers but without data assimilation was 1.71°C and 1.56°C, respectively (Figure 3). Data assimilation also removed a warming bias at 1 m and reduced the rate of warming through the spring and summer of 2019 at 8.0 m to more closely align model predictions with observed data (Figure 3). The three tuned GLM parameters were well-constrained by data assimilation (Supporting Information Figure 1) and their values varied following expected seasonal patterns.

**Figure 3.**
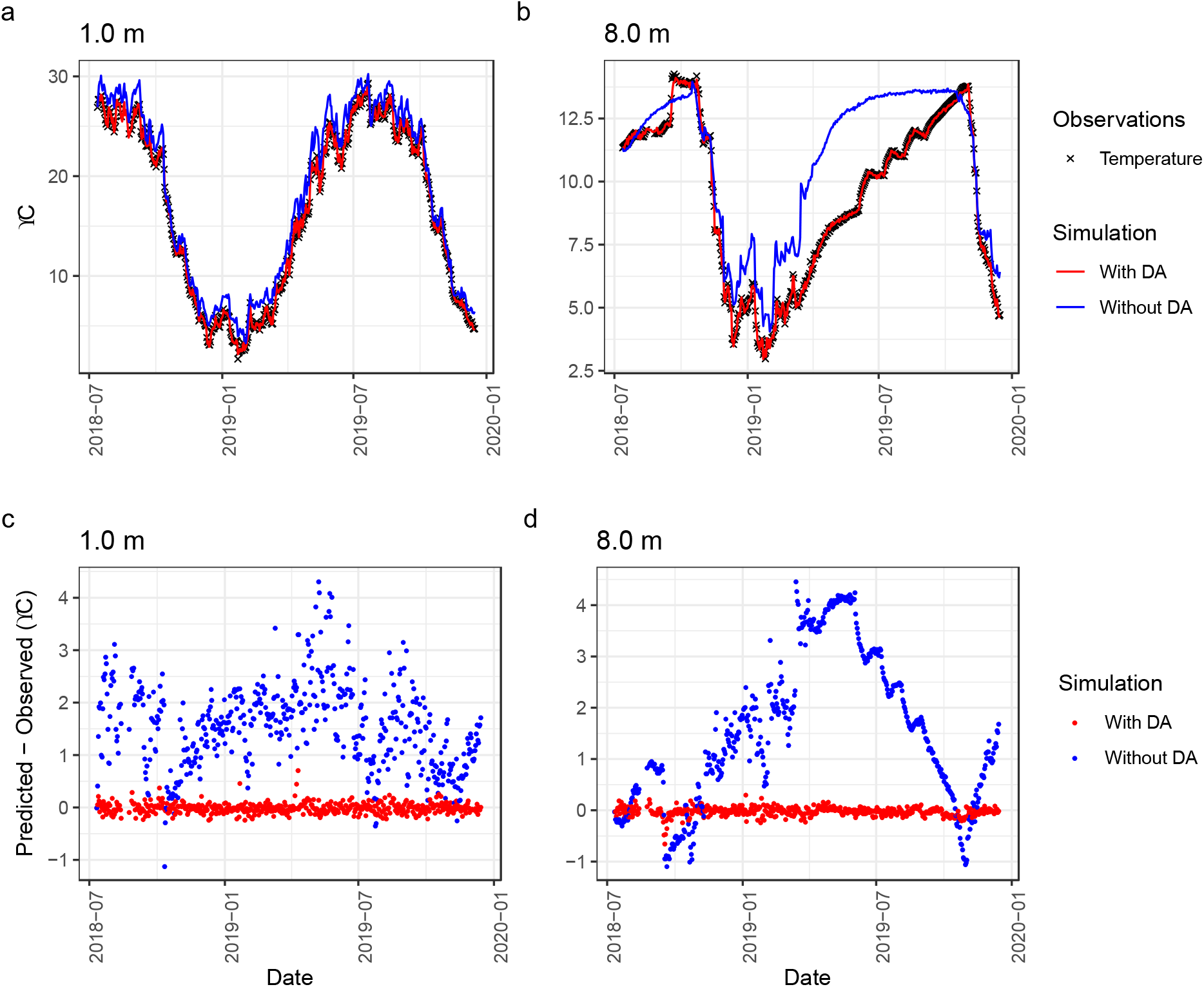
Mean predictions of water temperature at 1.0 (a) and 8.0 m (b) with (red lines) and without (blue lines) data assimilation of *in situ* observations of temperature measured at 10 depths. All simulations used observed meteorological and inflow drivers. Panels (c) and (d) show the bias in predictions at 1.0 and 8.0 m, respectively, with and without data assimilation.

### 3.3 Forecast performance

Every day between 28 August 2018 and 15 December 2019, the forecasting system generated 16-day forecasts of water temperature for the entire water column (see Figure 4 as an example). In general, forecast accuracy was high throughout the 16-day forecast horizon, with the mean bias in forecasted water temperatures within 0.30°C for 1.0 m and 0.22°C for 8.0 m of observed temperatures throughout the 475 daily forecasts, averaged across all forecast time horizons and ensemble members (Figure 5A; Table 1). RMSE was 0.22-0.40°C lower for 8.0 m forecasts than for 1.0 m forecasts aggregated across horizons and ensemble members, ranging from 0.30°C for 8.0 m at a 1-day horizon to 1.62°C for 1.0 m at a 16-day horizon (Table 1).

**Figure 4.**
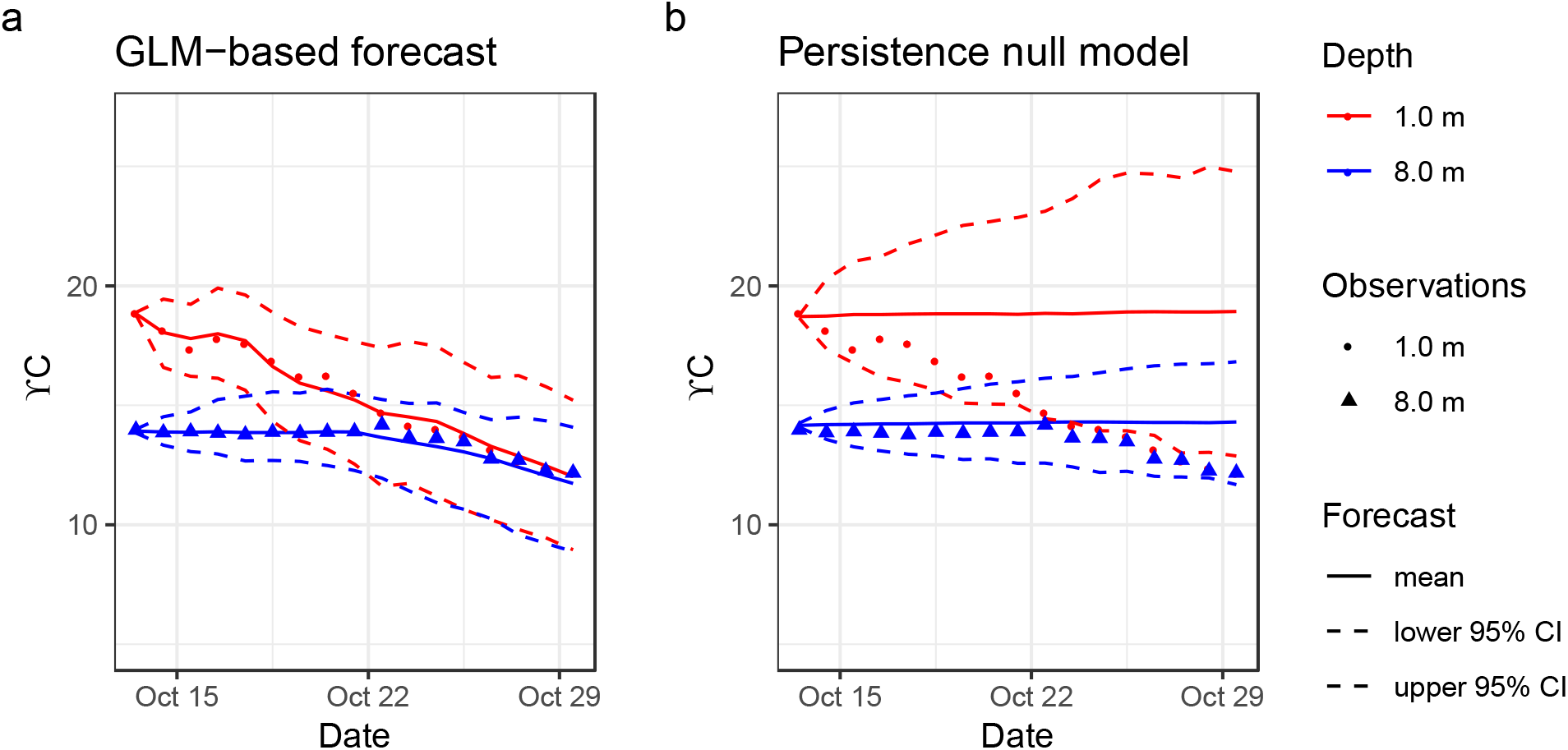
An example of a 16-day forecast (initiated on 13 October 2018) at the near-surface (1.0 m) and near-sediment (8.0 m) depths by the (a) General Lake Model and (b) the persistence null model. The forecast uncertainty in both panels is compared to the observations from the water temperature sensors at 1.0 and 8.0 m depths.

**Table 1.** Forecast evaluation metrics for the entire forecasting period (“all conditions”), during thermally-stratified conditions only, and during mixed conditions only. Metrics are calculated for all depths averaged together (“all depths”), at the near-surface (1.0 m), and at the near-sediments (8.0 m). RMSE (the root mean squared error between observations and forecasted temperatures); CRPS_forecast_ (Continuous Ranked Probability Score for the FLARE forecast); CRPS_null_ (CPRS for the persistence null forecast); CRPS_forecast_ Skill (skill score for the FLARE forecasting based on CRPS_forecast_ and CRPS_null_); bias (the difference between observations and forecasted temperatures); and confidence interval (CI) reliability (the percentage of observations within the 90% confidence intervals).

| Forecast time horizon | Metric | All conditions<br>(n = 475) |  |  | Stratified conditions<br>(n = 282) |  | Mixed conditions<br>(n = 193) |  |
| --- | --- | --- | --- | --- | --- | --- | --- | --- |
|  |  | 1.0 m | 8.0 m | All depths | 1.0 m | 8.0 m | 1.0 m | 8.0 m |
| 1 day | RMSE (°C) | 0.52 | 0.30 | 0.44 | 0.56 | 0.23 | 0.45 | 0.39 |
|  | CRPS<sub>forecast</sub> (°C) | 0.29 | 0.16 | 0.23 | 0.32 | 0.12 | 0.25 | 0.22 |
|  | CRPS<sub>null</sub> (°C) | 0.34 | 0.13 | 0.23 | 0.40 | 0.09 | 0.26 | 0.18 |
|  | CRPS<sub>forecast</sub> skill (unitless) | 0.16 | -0.26 | -0.07 | 0.21 | -0.3 | 0.04 | -0.23 |
|  | Bias (°C) | 0.07 | 0.04 | 0.03 | 0.07 | -0.03 | 0.09 | 0.15 |
|  | CI reliability (%) | 90 | 76 | 85 | 93 | 74 | 85 | 79 |
| 7 day | RMSE (°C) | 1.13 | 0.87 | 1.04 | 0.98 | 0.7 | 1.32 | 1.07 |
|  | CRPS<sub>forecast</sub> (°C) | 0.64 | 0.52 | 0.56 | 0.58 | 0.46 | 0.72 | 0.61 |
|  | CRPS<sub>null</sub> (°C) | 1.14 | 0.47 | 0.77 | 1.26 | 0.28 | 0.95 | 0.76 |
|  | CRPS<sub>forecast</sub> skill (unitless) | 0.44 | -0.1 | 0.20 | 0.54 | -0.66 | 0.24 | 0.20 |
|  | Bias (°C) | 0.04 | 0.00 | 0.05 | -0.16 | -0.27 | 0.34 | 0.39 |
|  | CI reliability (%) | 92 | 61 | 82 | 99 | 46 | 82 | 82 |
| 16 day | RMSE (°C) | 1.62 | 1.20 | 1.40 | 1.68 | 1.13 | 1.53 | 1.30 |
|  | CRPS<sub>forecast</sub> (°C) | 0.92 | 0.79 | 0.80 | 0.95 | 0.83 | 0.87 | 0.75 |
|  | CRPS<sub>null</sub> (°C) | 1.93 | 0.92 | 1.40 | 2.11 | 0.56 | 1.68 | 1.45 |
|  | CRPS<sub>forecast</sub> skill (unitless) | 0.53 | 0.14 | 0.39 | 0.55 | -0.48 | 0.48 | 0.49 |
|  | Bias (°C) | -0.03 | -0.06 | 0.03 | -0.45 | -0.56 | 0.57 | 0.66 |
|  | CI reliability (%) | 90 | 56 | 79 | 94 | 36 | 85 | 85 |

Forecast error as indicated by the CRPS_forecast_ increased as the forecast horizon increased, though it was notable that the CRPS_forecast_ was consistently better than the null persistence model (CRPS_null_) throughout the 16-day horizon for 1.0 m temperatures (Figure 5B). This resulted in a CRPS_forecast_ skill score for the 16-day time horizon of 0.53 when averaged across all forecasts. At 8.0 m depth, the null persistence model (CRPS_null_) performed better than the forecasts at the 1 and 7-day time horizons and only marginally worse at the 16-day time horizon. This resulted in a lower CRPS_forecast_ skill score of 0.14 at 8.0 m when averaged across all forecasts (Table 1).

**Figure 5.**
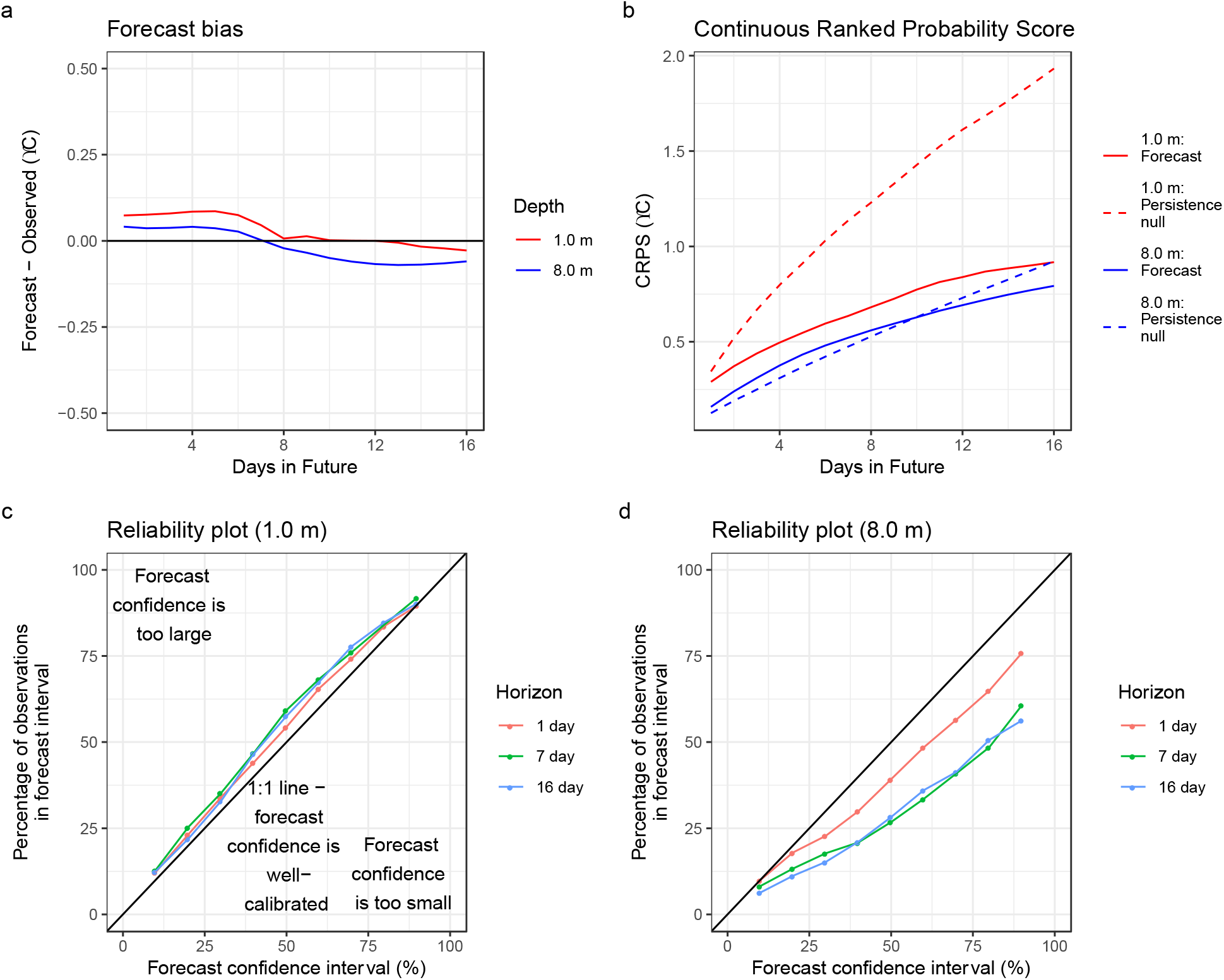
Water temperature forecasts for 1.0 m (red) and 8.0 m (blue) had low bias, high forecast skill (relative to a null persistence model), and well-calibrated confidence intervals. (a) The difference between the forecasted and observed water temperatures (bias) at 1.0 m and 8.0 m averaged across all forecasts generated during the 475-day forecasting period. (b) The Continuous Ranked Probability Score (CRPS, a metric for evaluating ensemble forecasting that is analogous to Mean Absolute Error) of the forecasts at the two depths averaged across the entire forecasting period in comparison to a null persistence model. The null model assumed that water temperature did not change over the 16-day forecast horizon. The percentage of (c) 1.0 m and (d) 8.0 m observations during the forecasting period that were within the forecast confidence interval that is specified on the x-axis. For example, the 90% forecast confidence interval encompasses the 5 th to 95th percentile of the 441 forecast ensemble members and ideally should contain 90% of water temperature observations. Values for 1, 7, and 16-day forecast horizons for the 1.0 m depth are shown.

Forecast confidence intervals at 1.0 m depth were well-calibrated (exactly 90% of the observations were in the 90% confidence intervals; Table 1) and generally fell along the 1:1 line between the percentage of observations within a specific forecast confidence interval (Figure 5C). Forecast confidence intervals were less well-calibrated at 8.0 m (56-76% of observations were within the 90% interval) and underestimated the uncertainty across all time horizons (Table 1; Figure 5D). Averaged across all depths, the forecast confidence intervals were smaller than expected (79-85% of observations fell within the 90% confidence intervals; Table 1).

Forecasting performance was generally similar between the thermally-stratified and mixed periods, with a few key exceptions. At 1.0 m, the CRPS forecast skill was consistently greater (up to 0.3 skill score) and bias was lower in stratified than mixed conditions, regardless of forecast time horizon (Table 1). In contrast, at 8.0 m, CRPS forecast skill was slightly greater at 1 and 7-day horizons but there was no difference in forecast skill between mixed and stratified conditions at a 16-day horizon (Table 1). Finally, the confidence intervals were better calibrated at 8.0 m for the mixed period (85% of observations in the 90% confidence interval at the 16-day time forecast horizon) than the stratified period (36% of observations in the 90% confidence interval; Table 1).

The forecasts successfully predicted the onset of fall turnover that occurred on 22 October 2018 up to 14 days in advance (Figure 6a). At 14 days prior to 22 October 2018, the predicted chance of turnover occurring on that day emerged above 50% for the first time (Figure 6a). Confidence that turnover would occur on 22 October increased eight days prior to turnover, when its predicted chance exceeded 75%. By 19 October (three days prior to turnover), the predicted chance of turnover on 22 October increased up to 81%, 31% higher than any of the other potential days.

**Figure 6.**
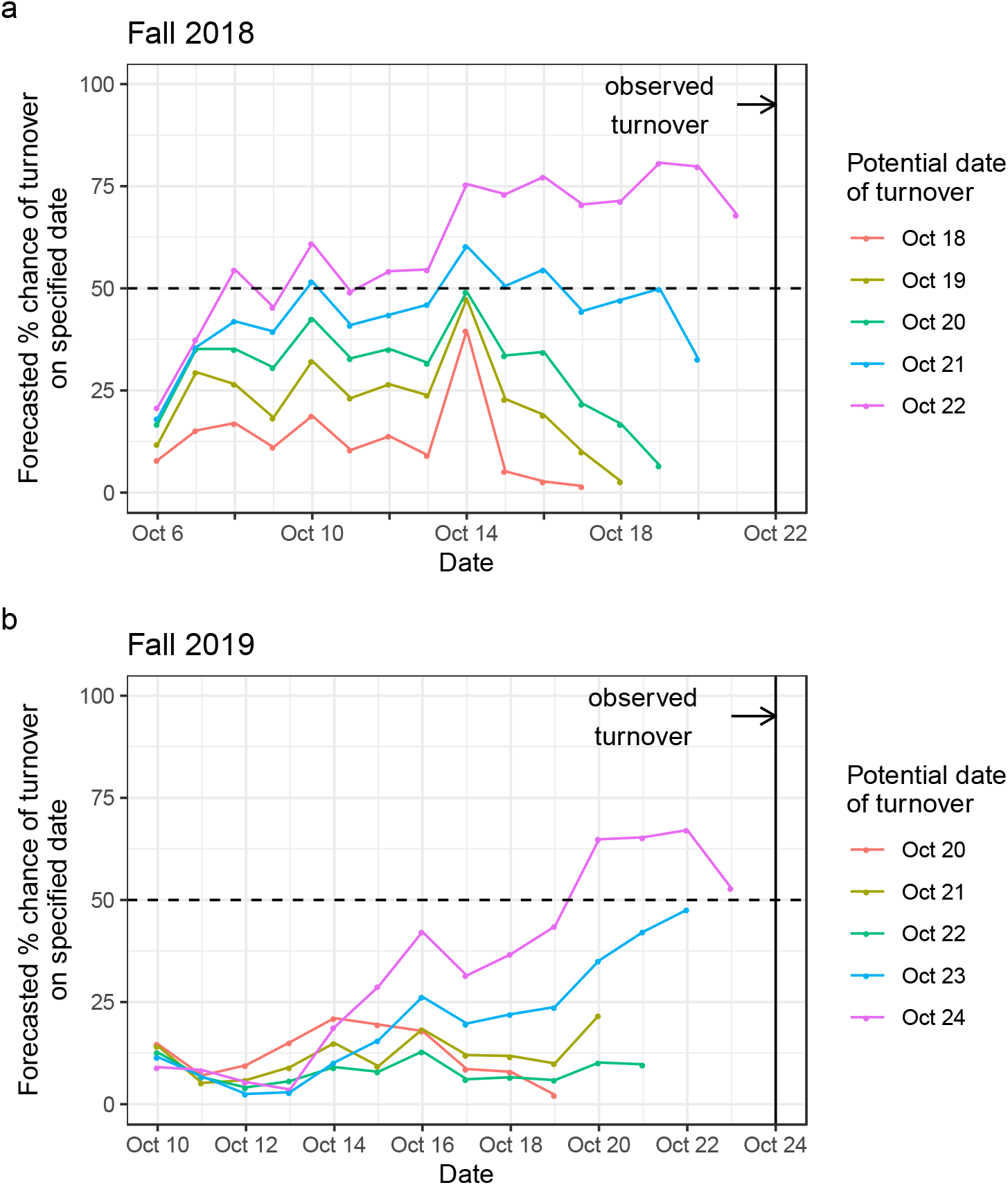
The forecasting system was able to anticipate the day of fall turnover (a) ~14 days in advance in 2018 and (b) ~4 days in advance in 2019 (as defined by when the percent chance first exceeded 50%). Turnover is defined as the first day in autumn when water temperatures at 1.0 and 8.0 m were <1°C at 12:00:00 UTC. Each panel shows how the percent chance of turnover for a particular forecasted day updated daily until turnover was observed.

The forecasts successfully predicted the onset of fall turnover that occurred on 24 October 2019 up to four days in advance (Figure 6b). The predicted chance of turnover occurring that day increased from 43% five days prior to turnover to 63% four days prior to turnover. The predicted chance of turnover occurring on 24 October remained above 50% as the day of turnover approached. However, the confidence in turnover date was lower in 2019 than 2018, potentially because the reservoir temporarily re-stratified immediately after the initial onset of mixed conditions in 2019 (Figure 2).

### 3.4 Uncertainty partitioning

Forecast uncertainty varied over time and depth. In the thermally-stratified period, the total forecast uncertainty was approximately four times higher for forecasts at 1.0 m than 8.0 m, with striking differences in the relative importance of different sources to total forecast uncertainty (Figure 7). For both depths, model process uncertainty was the most important contributor to total uncertainty over the first three days in a 16-day forecast horizon during stratified conditions. After three days, the contribution of meteorological driver uncertainty dominated at 1.0 m depth (>50% of uncertainty) while process uncertainty remained the dominant source of uncertainty at 8.0 m depth throughout the 16-day forecast horizon (>95% of uncertainty). At the near-surface, the statistical downscaling of the meteorological driver data was a more important uncertainty source than the uncertainty due to the NOAA meteorological forecast itself throughout the 16-day horizon. Parameters, inflow driver data, and initial conditions were not important uncertainty sources at either depth.

**Figure 7.**
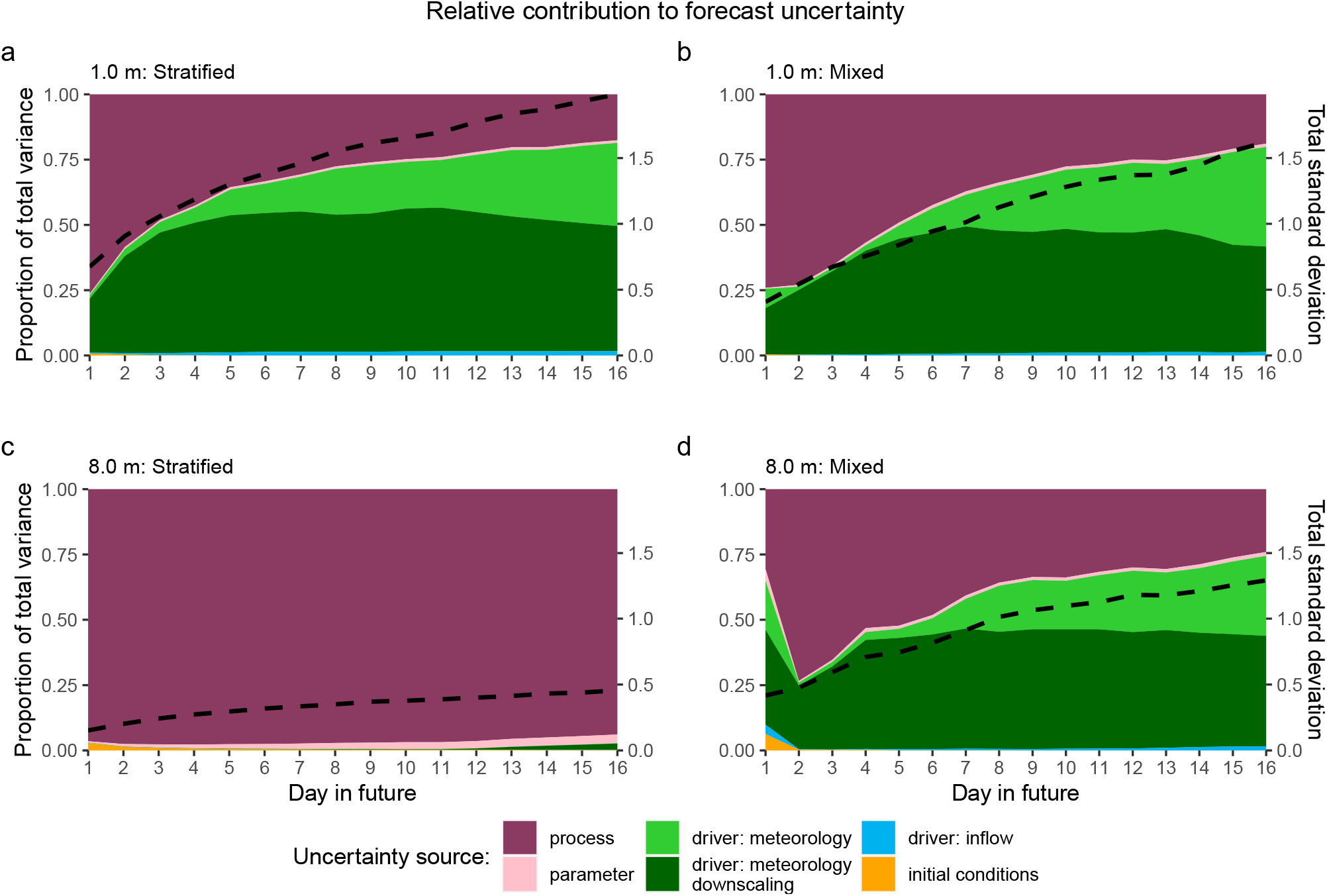
The relative contribution of the individual sources of uncertainty (left axis) to the total forecast uncertainty (right axis, black dashed line) varies through the 16-day forecast horizon and between stratified (left column) and mixed (right column) conditions at both 1.0 m depth (top row) and 8.0 m depth (bottom row). The relative contributions and total forecast variance for each day are averaged across 16 forecasts: N = 9 during summer stratification (a, c) and N = 7 during mixed conditions (b, d). Two depths are shown (1.0 m - a and b; 8.0 m - c and d).

In contrast to the thermally-stratified period, the total uncertainty and the relative contribution of the different sources were similar between the near-surface (1.0 m) and near-sediment (8.0 m) depths during mixed conditions (Figure 7). In the mixed period, process uncertainty was the most important source for the first five days of the forecast before the relative importance of the combined meteorological downscaling and forecasts dominated. Similar to the stratified period, parameters, inflow driver data, and initial conditions were minimal sources of uncertainty during mixed conditions.

## 4. Discussion

Overall, FLARE was able to forecast water temperature on average within 1.4°C (RMSE) at all depths of the reservoir over a 16-day horizon during a 475-day period that encompassed both stratified and mixed thermal conditions (Table 1). Importantly, the forecasting system was able to predict both observed temperatures and identify the date of fall turnover at least four, and up to 14, days in advance. In general, forecasting system performance was similar between stratified and mixed periods (Table 1), suggesting that the system is likely robust in a range of reservoir conditions, though additional forecasts are needed to provide a full assessment of FLARE performance. Generally, 1-D hydrodynamic models used for hindcasting aim to predict water temperature within an RMSE of 2°C [e.g., *Bruce et al.*, 2018], so the level of accuracy associated with FLARE future forecasts over 475 days at Falling Creek Reservoir exceeds expectations (RMSE = 1.4°C aggregated across all periods and all depths; Table 1).

We found that process uncertainty was the most important source of uncertainty early in the 16-day forecast but that meteorological driver data uncertainty dominated by the end of the forecasting period for all but near-sediment depths during the thermally-stratified period (Figure 7). This finding is intuitive, as the epilimnion is more sensitive to meteorological forcing than the hypolimnion, and during mixed conditions, both surface and near-sediment depths exhibit similar dynamics. We note that process uncertainty did not decline through the forecast horizon, rather, the total uncertainty grew while the relative importance of process uncertainty decreased. By definition, process uncertainty is not attributed to an individual component of a model but a model’s overall inability to represent complex interactions occurring in the natural environment [*Dietze*, 2017a]. Improvements to the GLM model structure, such as improved sediment heating formulations, could decrease this uncertainty source; however, there is likely an irreducible component of process uncertainty inherent to using a 1-D model to simulate a 3-D waterbody.

The contribution of the NOAA ensemble forecast uncertainty to total forecast uncertainty was overall less than the contribution of uncertainty from downscaling the coarse-scale NOAA forecast to the local site. This downscaling was based on relationships between the NOAA GEFS forecasted variables and the meteorological data measured at the reservoir (see Supporting Information C, D). This highlights that future work should focus on applying more advanced meteorological downscaling methods, such as neural networks [*Kumar et al.*, 2012], to build better relationships between the NOAA forecasts and the local meteorological station. Our uncertainty results are comparable to [*Dietze,* 2017b], who partitioned the uncertainty in forest carbon flux forecasts over 16-day horizons and similarly found that NOAA ensemble driver data uncertainty dominated total forecast uncertainty at the end of the 16-day horizon, though that study did not include NOAA forecast downscaling uncertainty. *Dietze* [2017b] also found that process uncertainty was more important than the meteorological driver data uncertainty early in the 16-day forecast horizon. Overall, while that study and ours are just two examples of forecast uncertainty partitioning, they provide insight from very different ecosystems (reservoir water temperatures vs. forest carbon fluxes) and together suggest that meteorological driver data are an important contributor of uncertainty in near-term iterative forecasts. Moreover, our study indicates that future forecasting applications should investigate the relative role of meteorological forecasts vs. their downscaling when quantifying total forecast uncertainty.

Throughout the forecasting period, inflow driver data, parameters, and initial conditions were minimal sources of uncertainty, regardless of forecast horizon. During the forecasting period, the reservoir exhibited a median water residence time of 145 days [*Carey et al.*, 2020c; *Gerling et al.*, 2014], so it was surprising that the inflow driver data did not contribute more to total uncertainty. However, the low residual error in our inflow forecast model (eqn. 2; R2 = 0.98) suggests that our simple first-order autoregressive model was able to successfully capture variability in inflow dynamics throughout the forecasting period. Inflow driver data may contribute more uncertainty for waterbodies with multiple inflows and higher discharge than we observed in our case study. Parameter uncertainty and initial conditions uncertainty were similarly minor, indicating that the data assimilation with state updating and automated parameter tuning that was built into the iterative forecasting cycle was able to constrain these sources of forecast uncertainty in our application.

FLARE was able to identify the onset of fall turnover ~14 days in advance in 2018 and approximately four days in advance in 2019. This determination was made by identifying the first day when the probability of turnover exceeded 50% (Figure 6). This difference in performance between years may be due to differing underlying hydrodynamic and meteorological drivers of fall turnover. In 2018, the surface waters of the reservoir had rapidly cooled (by ~8°C) in the 14 days prior to the onset of turnover in response to cold night air temperatures (down to 3°C) that continued after turnover. The day prior to turnover onset in 2018 coincided with high wind speeds, up to 9.8 m s-1 [*Carey et al.*, 2020a], which persisted after turnover and promoted continuous mixing once turnover started. In contrast, in 2019, the reservoir re-stratified later on the same day that the turnover criterion was first met due to warm daytime air temperatures (up to 18.7°C on 24 October) and wind speeds were approximately half of what was observed in 2018. The reservoir did not continuously exhibit isothermal conditions in 2019 until 2 November, eight days after the turnover criterion was initially met. The *McClure et al.* [2018] criterion of turnover was only met for 17 hours on 24 October 2019, making it extremely challenging for the forecasting system to correctly predict turnover on that particular day. Consequently, it is likely that using a more conservative fall turnover criterion (e.g., the temperature difference between 1 and 8 m depth must be <1oC and maintained for 48 consecutive hours, following *Woolway et al.* [2014]), would improve forecasting performance for future turnover forecasting applications.

We anticipate that FLARE may be of interest to managers of lakes and reservoirs where water temperature forecasts up to 16 days in advance can provide decision support. Many reservoirs, including FCR, have dynamic water extraction schedules, which are chosen based on water temperature depth profiles and the magnitude of thermal stratification [*Mi et al.*, 2019]. Knowing water temperature depth profiles *a priori* provides managers with valuable information for optimizing the temperature of downstream releases, water quality management, and *in situ* management prior to water treatment [*Caissie et al.*, 2016; *Huang et al.*, 2011; *Mi et al.*, 2019; *Pike et al.*, 2013; *Weber et al.*, 2017]. Given FCR’s small surface area and shallow depth, it would be expected that the forecasting system would be more accurate in larger, deeper lakes that are less sensitive to meteorological forcing. However, the difference in water temperature forecasts generated from the null persistence model and FLARE would likely be smaller in those bigger ecosystems.

Importantly, managers could use their current water temperature sensors for setting up and running FLARE to generate forecasts for their waterbody of interest, without needing expensive instrumentation or software. All software components of the FLARE system are opensource and the automated parameter tuning of the GLM model as part of the EnKF enables quick model setup. To deploy FLARE, a utility would need water temperature sensors and an inflow discharge sensor connected via a sensor gateway to the internet, meteorology data, and minimal knowledge of the waterbody’s bathymetry to set up boundary conditions in the GLM model. In comparison to many existing water temperature forecasting systems that rely on substantial sensor and satellite infrastructure for real-time forecasting [e.g., *Baracchini et al.*, 2020b; *Hague and Patterson,* 2014; *Ouellet-Proulx et al.*, 2017a; *Pike et al.*, 2013], FLARE provides a low-cost and open-source alternative with high forecasting performance.

While the FLARE forecasting system presented here was able to predict water temperature over the 16-day time horizon with low bias and an error substantially lower than a null persistence model, there are two important potential improvements. First, FLARE is using a physics-based, 1-D hydrodynamic model, so improvements to the physical parameterization and numerical solution would improve forecast skills, though we note that a 3-D model may be more time and compute-intensive. Forecasting approaches that combine physics-based models with machine learning [e.g., *Jia et al.*, 2019; *Read et al.*, 2019] could reduce forecast uncertainty by leveraging the power of mechanistic models and machine learning methods. However, machine learning-based methods must be able to fully quantify forecast uncertainty to be comparable to our process model-based approach [see *Daw et al.*, 2020]. Second, our parameter uncertainty may be underestimated because we only included three parameters in the EnKF data assimilation, though we note that our sensitivity analysis determined those three parameters to be the most important for water temperature predictions within the GLM (Supporting Information A). While the ability to estimate parameters using EnKF is well-established, the EnKF method is not specifically designed to estimate parameter distributions like Bayesian Monte-Carlo Markov Chain methods [*Dietze,* 2017a]. A current limitation to implementing a Bayesian Monte-Carlo Markov Chain approach is the computation time to execute the GLM simulation within a daily iterative forecasting cycle. Future work that uses emulators of GLM may be able to speed computation and allow for more robust estimation of the joint distribution model of parameters that represent both prior knowledge and observed data [e.g., *Fer et al.*, 2018].

We note four limitations of our application of FLARE at FCR. First, our implementation included only one inflow, in which we used a simple first order auto-regressive model to forecast future inflow discharge and water temperature. Further work is needed to evaluate FLARE in waterbodies with more inflows. For those waterbodies, FLARE could be linked to a watershed hydrology model to mechanistically link the precipitation and temperature forecasts to inflow driver data. Second, we set outflow equal to inflow to maintain a constant water level based on observations, which does not occur in many lakes and reservoirs. Future development of FLARE could explicitly assimilate observations of water level to constrain forecasts in more dynamic systems. Third, our data assimilation in FLARE used high-frequency sensors measuring water temperature at 1-meter depth intervals at 10-minute resolution, an observational capacity at FCR that may not be available in all waterbodies. Future evaluation of FLARE should focus on determining how forecast quality degrades as the vertical and temporal resolution of observations are reduced to reflect other common temperature monitoring approaches (e.g., weekly temperature depth profile measurements with a hand-held sensor, minute to hourly temporal observation at only one depth, remote sensing measurements of surface temperature, etc.). Finally, our application was limited to a period with minimal ice cover; further work is needed to determine FLARE performance for FCR during longer ice-covered periods and waterbodies that experience ice for many consecutive months.

Overall, our study demonstrates the utility of the daily iterative forecast workflow (Figure 1) for forecasting lake and reservoir water temperatures with fully-specified uncertainties that can be applied to other waterbodies. FLARE builds the foundation for future water quality data assimilation and forecasting because multiple ecosystem models can easily be coupled to the GLM hydrodynamic model [e.g., *Hipsey et al.*, 2013; *Snortheim et al.*, 2017], enabling predictions of dissolved oxygen, algal blooms, and biogeochemical cycling with uncertainty. Importantly, FLARE provides a method for partitioning uncertainty in forecasts that identifies how to prioritize future research to increase confidence in forecasts. Given the pressing need for tools to anticipate the increasing variability of freshwater ecosystems, near-term iterative forecasting systems such as FLARE provide the ability to anticipate future change for stakeholders, managers, and policy-makers.

## Data and Code Availability

All data used in this study are available in the Environmental Data Initiative repository [*Carey,* 2019; *Carey et al.*, 2020a; *Carey et al.*, 2020b; *Carey et al.*, 2020c] and through NOAA forecast archives (https://www.ncdc.noaa.gov/data-access/model-data/model-datasets/global-ensemble-forecast-system-gefs). Code for FLARE can be found at: https://github.com/CareyLabVT/FLARE/releases/tag/v1.1 Code for the sensor gateways can be found at: https://github.com/CareyLabVT/SCC_gateway

## Author Contributions

RQT, CCC, and RJF co-developed the forecasting system framework and workflow. RQT developed the data assimilation system, RJF and VD developed the cyberinfrastructure, CCC and BJB deployed the water quality sensors, BJB and VD maintained the sensor and data workflows, and LKP and RQT developed and applied the meteorology downscaling technique. CCC and RQT wrote the manuscript; all authors provided feedback and approved the final version.

