## Supporting Information for "A near-term iterative forecasting system successfully predicts reservoir hydrodynamics and partitions uncertainty in real time"

**Pages:** 21

**Text:** 4

**Tables:** 3

**Figures:** 1

**Supporting Information A. Detailed description of the data assimilation methods used in FLARE.**

The data assimilation in FLARE uses the ensemble Kalman Filter (EnKF) with state augmentation to set the initial variables for the states (initial conditions) and parameters for each 16-day forecast. The EnKF updates predictions of water temperature at specified depths (the states) and calibrates parameters using observations of water temperature at multiple depths (following the methods of *Zhang et al.* [2017]). The EnKF is initialized at the beginning of forecasting and data assimilation (i.e., $t$ = 0 on the day the automated forecasting system is deployed) with a set of $M$ ensemble members, in which each ensemble member $i$ has a vector of modeled water temperatures at $K$ depths at the *0*th time ($x_{t=0}^{i}$) and a vector of $P$ parameters ($\alpha_{t=0}^{i}$). The values of $x_{0}^{i}$ are initialized with observed sensor temperatures and linear interpolation is used to initialize the modeled depths that did not have observations.

For this application, $P$ was three because three General Lake Model (GLM) parameters were calibrated for Falling Creek Reservoir: SW_factor, zone1temp, and zone2temp. These parameters were chosen based on a global sensitivity analysis of all GLM parameters. We used the elementary effects sensitivity analysis method outlined by *Morris* [1991], in which we randomly sampled distributions of each GLM parameter in 10,000 simulations and calculated the mean and standard deviation of the distribution of elementary effects for each parameter. Following *Morris* [1991], the mean of a parameter's elementary effects is a metric of the parameter's overall influence on model output, and its standard deviation is a metric of its effect on other parameters. Next, we applied K-means cluster analysis to the mean and standard deviations of each parameter's elementary effects distributions. SW_factor, LW_factor, zone1temp, and zone2temp emerged in a unique cluster with the highest mean and standard deviation of elementary effects. Because SW_factor and LW_factor are tightly coupled in the GLM, we included only SW_factor in the calibration, as it had a slightly higher mean elementary effects than LW_factor. Finally, $\alpha_0^i$ was initialized using a random draw for each ensemble from a parameter-specific uniform distribution (ranges for SW_factor: 0.5 – 1.5; zone1temp: 10-20$_{o}$C; and zone2temp: 10-20$_{o}$C). We used uniform distributions because we did not have any data to justify using any other type of distribution; the ranges were set to be large enough to include any reasonable value for the parameters, following *Hipsey et al.* [2019].

Every day following the deployment of the forecasting system, the observed meteorology from the previous 24 hours is downloaded from a GitHub repository to a local server and automatically processed to generate a matrix of hourly meteorological inputs for GLM. This matrix was combined with the other model driver data (inflow discharge rate and inflow water temperature) to create a driver matrix $(D_t^i)$ for each ensemble member, where *t* is the number of days since forecasting system deployment. The GLM input drivers during this 24-hour historical period were the same for all ensemble members.

The vector of modeled water temperature for each depth from the previous day $(x_{t-1}^i)$, the parameter vector $(\alpha_{t-1}^{i-})$, and the last 24 hours of driver data $(D_t^i)$ were used to initialize and run a 1-day simulation of the GLM for each ensemble member, $G\left(x_{t-1}^i, \alpha_{t-1}^i, D_t^i\right)$. Process uncertainty was added to the water temperature predictions from the GLM following eqn. SI.1 to create predictions of water temperature with process uncertainty for each depth:

$$x_t^{i-} = G\left(x_{t-1}^i, \alpha_{t-1}^i, D_t^i\right) + \varepsilon \text{ (eqn. SI.1)}$$

where $x_t^{i-}$ is the $K \times 1$ vector of predicted water temperatures at the modeled depths for the *i*th ensemble member at time *t*. $\varepsilon$ (a $K \times 1$ vector) is a random draw from a multivariate normal distribution with a mean of 0 and an covariance matrix $\Sigma_t$ at time *t*.

The $\Sigma_t$ matrix evolves after the deployment of the forecasting system through data assimilation, as the model predictions prior to updating $(x_t^{i-})$ improve or degrade over time. This allows for the process uncertainty to reflect the performance of model predictions over a specified time period (a 30-day window in our application for Falling Creek Reservoir). For the first 30 days after forecasting system deployment, the $\Sigma_t$ is a diagonal matrix with a constant variance for all depths (0.5°C) that does not evolve over time. After the first 30 days, a 30-day running covariance matrix at the observed depths $(\Sigma_t^*)$ is calculated as the residual of the predictions prior to updating, following eqn. SI.2:

$$\Sigma_t^* = \frac{1}{V}\sum_{l=t}^{V}(\overline{x_{t-l}} - y_{t-l})(\overline{x_{t-l}} - y_{t-l}) \quad \text{(eqn. SI.2)}$$

where $\Sigma_t^*$ is used to calculate $\Sigma_t$ by linearly interpolating the variances and covariances between depths in $\Sigma_t^*$. In eqn. SI.2, $V$ is the number of previous days included in the covariance matrix (here, 30). A 30-day window was chosen to balance two needs: 1) that enough days with observations were included in the analysis of the residuals to reduce spurious covariances in the covariance matrix, and 2) that the residuals reflect recent model performance.

If data are not available to update the model states due to missing sensor data, the states are not updated and $x_t^i = x_t^{i-}$. Otherwise, the covariance among states in the ensemble members $(C_{xx})$ is calculated using eqn. SI.3:

$$C_{xx} = \frac{1}{M-1}\sum_{i=1}^{M}(x_t^{i-} - \overline{x_t})(x_t^{i-} - \overline{x_t}) \quad \text{(eqn. SI.3)}$$

where $\overline{x_t}$ is the mean temperature at each modeled depth across ensemble members. The $C_{xx}$ matrix represents the estimated model error.

The covariance among parameters and states in the ensemble members $(C_{\alpha x})$ is similarly estimated using the relationship between parameters and model predictions following eqn. SI.4:

$$C_{\alpha x} = \frac{1}{M-1}\sum_{i=1}^{M}(\alpha_{t-1}^{i-} - \overline{\alpha_{t-1}})(x_t^{i-} - \overline{x_t}) \text{ (eqn. SI.4)}$$

where $\overline{\alpha_{t-1}}$ is the mean for the parameter vector across ensemble members. In order to prevent filter divergence that is caused by artificially small parameter variance [*Dietze*, 2017a], we apply ensemble inflation to the parameters prior to calculating $C_{\alpha x}$ in eqn SI.4, similar to the approaches used by *Zhang et al.* [2017] and *Page et al.* [2018]. The ensemble inflation is determined for each parameter using:

$$\alpha_{j,t}^{i-} = \overline{\alpha_{j,t-1}} + IF\left(\alpha_{j,t-1}^{i} - \overline{\alpha_{j,t-1}}\right) \text{ (eqn. SI.5)}$$

where *j* is the parameter index and *IF* is the variance inflation factor (here, we set *IF* = 0.02) and $\overline{\alpha_{j,t-1}}$ is the ensemble mean for parameter *j* at time step *t*-1.

Next, to quantify uncertainty in the observations, normally-distributed noise is added to the vector of observations at time t $(y_t)$ using the observation covariance matrix $(R)$ (eqn. SI.6):

$$\hat{y}_t^i = y_t + \varepsilon \text{ (eqn. SI.6)}$$

where $\hat{y}_t^i$ is the vector of observations and $\varepsilon$ is a random draw from a multivariate normal distribution with a mean of 0 and a covariance matrix R. In our application, the observational uncertainty is equal for all depths and not correlated among depths, and thus the $R$ matrix is diagonal.

The model states (i.e., the water temperatures at specific depths) and parameter updating using the observations first requires calculating the Kalman gain for the states $(K_x)$ and parameters $(K_\alpha)$ following eqn. SI.7:

$$\begin{bmatrix} K_x \\ K_\alpha \end{bmatrix} = \begin{bmatrix} C_{xx}H^T(HC_{xx}H^T + R)^{-1} \\ C_{\alpha x}H^T(HC_{xx}H^T + R)^{-1} \end{bmatrix} \text{ (eqn. SI.7)}$$

where *H* is a matrix in which each row corresponds to a depth with an observation and each column represents each of the modeled depths. The columns that correspond to the depths with temperature sensor observations have a value of 1 while all other columns have a value of 0. Each row only has a single 1. *T* represents the transpose of the *H* matrix.

The Kalman gain represents the proportional adjustment of the GLM model output based on the difference between the model predictions of water temperature and the sensor observations. The Kalman gain also updates depths without sensor observations when they are correlated with observed depths in the model predictions in $C_{xx}$. Similarly, the parameters are updated based on their correlation with the updated states. Following eqn. SI.8, the state gain $K_x(\hat{y}_t^i - Hx_t^{i-})$ and parameter gain $K_\alpha(\hat{y}_t^i - Hx_t^{i-})$ are added to the corrupted states ($x_t^{i-}$) and variance-inflated parameters ($\alpha_t^{i-}$):

$$\begin{bmatrix} x_t^i \\ \alpha_t^i \end{bmatrix} = \begin{bmatrix} x_t^{i-} \\ \alpha_t^{i-} \end{bmatrix} + \begin{bmatrix} K_x(\hat{y}_t^i - Hx_t^{i-}) \\ K_\alpha(\hat{y}_t^i - Hx_t^{i-}) \end{bmatrix} \text{(eqn. SI.8)}$$

**Supporting Information B. General Lake Model configuration namelist file.**

See [*Hipsey et al.*, 2019] for more information on the following General Lake Model configuration file (glm3.nml) for Falling Creek Reservoir (FCR), which describes the physical characteristics of the reservoir, parameters, and simulation setup.

```
&glm_setup
  sim_name = 'FCR'
  max_layers = 500
  min_layer_vol = 0.025
  min_layer_thick = 0.15
  max_layer_thick = 0.33
  non_avg = .true.
/
&light
  light_mode = 0
  n_bands = 4
  light_extc = 1.0, 0.5, 2.0, 4.0
  energy_frac = 0.51, 0.45, 0.035, 0.005
  Benthic_Imin = 10
  Kw = 0.87
/
&mixing
  coef_mix_conv = 0.2
  coef_wind_stir = 0.23
  coef_mix_shear = 0.3
  coef_mix_turb = 0.51
  coef_mix_KH = 0.30
  coef_mix_hyp = 0.50
  deep_mixing = 2
/
&morphometry
  lake_name = 'FallingCreek'
  latitude = 37.30768
  longitude = -79.83707
  bsn_len = 711.699
  bsn_wid = 226.03
  bsn_vals = 31
  H = 497.683, 497.983, 498.283, 498.683, 498.983, 499.283, 499.583, 499.883, 500.183,
500.483, 500.783, 501.083, 501.383, 501.683, 501.983, 502.283, 502.583, 502.883, 503.183,
```

```
503.483, 503.783, 504.083, 504.383, 504.683, 505.083, 505.383, 505.683, 505.983, 506.283,
506.583, 506.983
   A = 0, 61.408883, 494.615572, 1201.23579, 2179.597283, 3239.620513, 4358.358439,
5637.911458, 6929.077352, 8228.697419, 9469.324081, 10811.30792, 12399.67051,
14484.22802, 16834.20941, 19631.05422, 22583.1399, 25790.70893, 28442.99667,
31155.95008, 36269.3312, 42851.13714, 51179.89109, 59666.85885, 68146.39437,
76424.14457, 85430.25429, 95068.47603, 103030.4489, 111302.1604, 119880.9164
/
&time
   timefmt = 2
   start = '2018-04-19 12:00'
   stop = '2018-04-20 12:00'
   dt = 1800
   num_days = 1
   timezone = -5
/
&init_profiles
   num_depths = 10
   lake_depth = 9.4
   the_depths = 0.1, 1, 2, 3, 4, 5, 6, 7, 8, 9
   the_temps = 19.795, 19.914, 20.235, 20.166, 19.705, 17.53, 17.631, 17.599, 17.951, 18.245
   the_sals = 0, 0, 0, 0, 0, 0, 0, 0, 0, 0
   snow_thickness = 0.0
   white_ice_thickness = 0.0
   blue_ice_thickness = 0.0
   avg_surf_temp = 6.0
   restart_variables = 0, 0, 0, 0, 0, 0, 0, 0, 0, 0, 0, 0, 0, 0, 0, 0, 0
/
&meteorology
   met_sw = .true.
   lw_type = 'LW_IN'
   rain_sw = .false.
   atm_stab = 0
   catchrain = .false.
   rad_mode = 1
   albedo_mode = 1
   cloud_mode = 4
   meteo_fl =  !defined in FLARE!
   subdaily = .true.
   wind_factor = 1.0
```

```
  sw_factor = !defined in FLARE!
  lw_factor = 1.0
  at_factor = 1
  rh_factor = 1
  rain_factor = 1
  cd = 0.0013
  ce = 0.0013
  ch = 0.0013
  time_fmt = 'YYYY-MM-DD hh:mm:ss'
/
&inflow
  num_inflows = 1
  names_of_strms = 'weir'
  subm_flag = .true.
  strm_hf_angle = 55
  strmbd_slope = 0.05
  strmbd_drag = 9.00
  inflow_factor = 1.0
  inflow_fl = !defined in FLARE!
  inflow_varnum = 3
  inflow_vars = 'FLOW', 'TEMP', 'SALT'
/
&outflow
  num_outlet = 1
  flt_off_sw = .false.
  outl_elvs = 506.9
  bsn_len_outl = 711.699
  bsn_wid_outl = 226.03
  outflow_fl = 'outflow_file1.csv'
  outflow_factor = 1.0
/
&snowice
  snow_albedo_factor = 1
  snow_rho_max = 500
  snow_rho_min = 100
/
&sediment
  benthic_mode = 2
  sed_heat_model = 1
  n_zones = 2
```

```
   zone_heights = 5, 9.5
   sed_heat_Ksoil = 1.2, 1.2
   sed_temp_depth = 0.5, 0.5
   sed_temp_mean = !defined in FLARE!, !defined in FLARE!
   sed_temp_amplitude = 0, 0
   sed_temp_peak_doy = 257, 251
/
```

**Supporting Information C. Description of how the NOAA GEFS forecasts were spatially- and temporally downscaled.**

The overarching goal of the spatial and temporal downscaling was to adjust the 1º x 1º spatial resolution and 6-hour temporal resolution NOAA GEFS forecasts to represent the reservoir's local meteorological conditions at a 1-hour temporal resolution.

First, we used historical GEFS forecasts and 1-minute scale observational data measured at the reservoir from 12 July 2018 – 11 July 2019 as the "training data" for the spatial downscaling [*Carey et al.*, 2020a]. We aggregated both the NOAA GEFS and the observed meteorology to the daily scale by averaging all observations (except for precipitation, which was summed) and matched the data by date. In this training dataset, we only used the first day of each historical 16-day NOAA GEFS forecast because NOAA GEFS ensemble members were most similar to each other 1 day in the future and represent any consistent offsets between the 1º × 1º forecast and the local conditions.

To spatially-downscale temperature, relative humidity, wind speed, shortwave radiation, and longwave radiation, we estimated the linear relationship between the daily observation and forecast data in the training dataset (Supporting Information Table 2). We then applied this linear model to each day of the 16-day forecast. We set downscaled values for each variable that were less than zero to zero and values of relative humidity greater than 100 to 100. This resulted in a spatially-downscaled NOAA GEFS forecast product at the daily time scale.

To temporally-downscale the spatially-downscaled temperature, relative humidity, and wind speed forecasts from the daily to 1-hour resolution required by GLM, we first used the difference between the pre-spatially downscaled NOAA GEFS 6-hour forecast and its daily mean to convert the daily spatially-downscaled forecast to its original 6-hour resolution. We used a monotone Hermite spline method to obtain hourly values from the 6-hour values. Before applying the spline method within the first 6-hour period, we used the observed meteorology as the 0-hour variable and the downscaled forecast as the 6-hour value. This allowed for a smooth transition between the observed meteorology used in data assimilation and the downscaled forecast.

To temporally-downscale shortwave radiation from the spatially-downscaled daily resolution to 1-hour resolution, we calculated the potential top-of-atmosphere solar radiation for each hour to determine a scaling factor between hourly shortwave radiation and the mean daily potential shortwave radiation [following the solar_geom.R function in *Dietze*, 2017b]. We used this ratio to convert the daily downscaled shortwave radiation to the 1-hour resolution.

To temporally-downscale longwave radiation from the spatially-downscaled daily resolution to 1-hour resolution, we first used the relative difference between the pre-spatially downscaled NOAA GEFS 6-hour forecast and its daily mean to convert the daily spatially-downscaled forecast to its original 6-hour resolution. We then applied the 6-hour mean value to each hour within that time window.

Precipitation was only spatially-downscaled. We first calculated the ratio of the forecasted precipitation to observed precipitation in the training data. Then, we multiplied each NOAA GEFS 6-hourly forecasts of precipitation by this ratio and assumed constant precipitation for each hour within the 6-hour interval.

Finally, we represented uncertainty in the downscaling process by adding random noise to each spatial-downscaled daily forecast prior to temporally downscaling. First, we calculated the residuals between the observed meteorology and the downscaled NOAA GEFS forecast at the daily resolution for temperature, relative humidity, wind speed, shortwave radiation, and longwave radiation (as described below). This resulted in a set of residuals for each variable (except precipitation) within each hour. Second, we used the residuals to determine the covariance of residuals among variables across all hours in the training dataset (Supporting Information Table 3). Third, to add noise to each day of a 16-day forecast, we used this covariance to draw random values for each variable from a multivariate normal distribution that was centered at the downscaled values. Finally, the daily downscaled meteorological variables with the random noise added were temporal-downscaled as described above. In total, we generated 21 random draws from the downscaling uncertainty for each of the 21 downscaled NOAA GEFS ensembles on each day in the 16-day forecast horizon.

**Supporting Information D: A description of the sensor array at the reservoir and wireless data transmission methods**.

We measured the water temperature profile in Falling Creek Reservoir on 1-m depth intervals from the surface (0.1 m) to just above the sediments at 9 m at the deepest site of the reservoir with NexSens T-Node FR thermistors (NexSens Technology, Inc.; Fairborn, Ohio, USA; [*Carey et al.*, 2020b]). Thus, we had 10-minute sensor observations for 0.1 m, 1 m, 2 m, 3 m, 4 m, 5 m, 6 m, 7 m, 8 m, and 9 m beginning in July 2018. We were limited to forecasting from July 2018 to the present because the temperature sensors were not deployed in the reservoir prior to this period. The thermistor string was factory-calibrated and verified against a NIST-traceable thermistor to meet measurement accuracy of ±0.075$_{o}$C. These temperature observations were supplemented by a YSI EXO2 sonde temperature sensor collecting 10-minute observations at 1.6 m depth (YSI Inc., Yellow Springs, Ohio, USA). This sensor was factory-calibrated and had an accuracy of ±0.01$_{o}$C [*Carey et al.*, 2020b].

A Campbell Scientific (Logan, Utah, USA) research-grade meteorological station was deployed on the dam of the reservoir measured shortwave radiation, longwave radiation, air temperature, relative humidity, rainfall, wind speed, and barometric pressure in 2015 [*Carey et al.*, 2020a]. These meteorological variables were measured every minute and then downsampled (temperature, wind speed, humidity), averaged (shortwave and longwave), or summed (precipitation) to the hourly scale to serve as driver data for the GLM model (Supporting Information Table 1).

We measured the inflow discharge rate of the primary tributary entering into FCR from a pressure transducer deployed through a weir, which recorded the water temperature and water level every 15 minutes [*Carey et al.*, 2020c]. From 11 July 2018 - 21 April 2019, we measured water level and water temperature with an INW Aquistar PT2X pressure sensor (INW, Kirkland, Washington, USA), and from 22 April 2019 - 31 December 2019, we used a Campbell Scientific CS451 pressure transducer. The Campbell Scientific pressure transducer data were corrected via an offset to ensure there was no disruption in the time series from changing sensors. We used the water level to calculate the mean daily discharge rate following [*Gerling et al.*, 2014] and set the outflow discharge rate to the inflow discharge rate, as the reservoir was maintained at a constant water level through the study.

The water temperature, meteorological, and inflow sensor data were staged on Campbell Scientific data loggers on-site as measurements were retrieved, and transmitted daily to cloud storage. The sensor gateway attached to the Campbell Scientific data loggers ran the Ubuntu Linux software distribution, as well as software applications and scripts that were developed to perform data transfer and management functions including: 1) retrieve data from the logger using Campbell Scientific interfaces, 2) check cellular modem connectivity and reset modules as needed; and 3) reliably upload sensor data updates to appropriate Git repositories on cloud storage using the Git client. Data were structured as a time series, with measurements appended as lines to a comma-separated values (CSV) file. Data transfers used Git (https://git-scm.com), an open-source distributed version control system, for efficient and reliable updates with minimum bandwidth usage, such that only the data collected since the last successful transfer were sent from the gateway to the cloud server. The gateway also ran a virtual private network (VPN) open-source software, IPOP (IP-over-P2P) to provide authentication and encryption [*Ganguly et al.*, 2006], thereby providing a secure data transfer. All code used for configuring and running the sensor gateways can be found at: https://github.com/CareyLabVT/SCC_gateway

**Supporting Information Table 1**. Meteorological sensors deployed on the dam at the Falling Creek Reservoir as part of a research-grade Campbell Scientific weather station that collected driver data for the General Lake Model.

| Sensors deployed at the reservoir | Meteorological variables measured | Measurement precision |
|---|---|---|
| Rotronic Hydroclip2 HC2S3-L Temperature and Relative Humidity Probe with RM Young 10 plate Solar Radiation Shield | Air Temperature at 2 m | -50 - 100℃ ± 0.1℃ |
| | Relative Humidity at 2 m | 0 - 100% ± 1.3% |
| RM Young 05103-L Wind Monitor | Wind Speed at 4 m | 0 - 100 m/s ± 0.3 m/s |
| Hukseflux NR01 4-component Net Radiometer | Surface Downward Shortwave Radiation Flux | 0 - 2000 W/m² ± 10% |
| | Surface Downward Longwave Radiation Flux | 0 - 1000 W/m² ± 10% |

**Supporting Information Table 2**. The slope, intercept, and $R^2$ for the relationship between the first day of each NOAA GEFS forecast for the grid cell that contains Falling Creek Reservoir and the observed meteorology from the on-site weather station, as described in Supporting Information B.

| | Slope | Intercept | $R^2$ |
|---|---|---|---|
| Air temperature | 0.99 | 4.68 | 0.98 |
| Relative humidity | 1.1 | -14.88 | 0.63 |
| Wind speed | 0.47 | 0.84 | 0.46 |
| Shortwave radiation | 0.82 | 0.68 | 0.87 |
| Longwave radiation | 0.99 | 32.51 | 0.95 |

**Supporting Information Table 3.** Covariance matrix describing the relationships among residuals from the observed meteorology and downscaled NOAA GEFS forecasts (see Supporting Information B).

| | Air temperature | Wind speed | Relative humidity | Shortwave radiation | Longwave radiation | Rain |
|---|---|---|---|---|---|---|
| Air temperature | 1.75 | 0.06 | -4.32 | 5.30 | 4.24 | 0 |
| Wind speed | 0.06 | 0.27 | -0.48 | 4.33 | -1.56 | 0 |
| Relative humidity | -4.32 | -0.48 | 107.42 | -68.39 | 20.05 | 0 |
| Shortwave radiation | 5.30 | 4.33 | -68.39 | 987.84 | -163.35 | 0 |
| Longwave radiation | 4.24 | -1.56 | 20.05 | -163.35 | 153.71 | 0 |
| Rain | 0 | 0 | 0 | 0 | 0 | 0 |

**Supporting Information Figure 1**. Time series for the three calibrated parameters: a) shortwave factor, b) mean zone 1 (hypolimnetic) sediment temperature, and c) mean zone 2 (epilimnetic) sediment temperature from each ensemble over the analysis period.

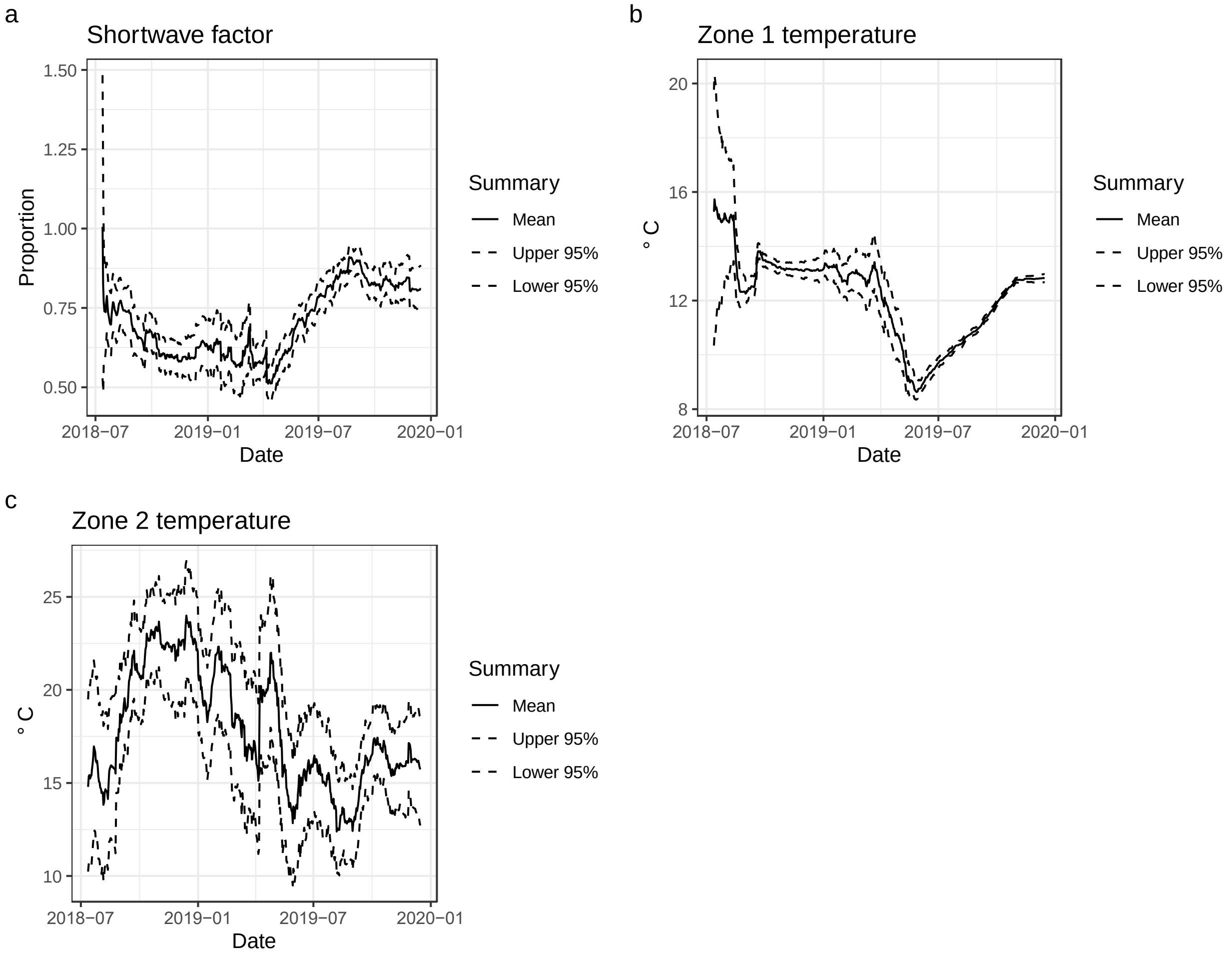
